# An adaptive pre-DNA-damage-response protects genome integrity

**DOI:** 10.1101/2020.05.13.092460

**Authors:** Sandrine Ragu, Gabriel Matos-Rodrigues, Nathalie Droin, Aurélia Barascu, Sylvain Caillat, Gabriella Zarkovic, Capucine Siberchicot, Elodie Dardillac, Camille Gelot, Juan Pablo Radicella, Alexander A. Ishchenko, Jean-Luc Ravanat, Eric Solary, Bernard S. Lopez

**Author notes:** Correspondence to: Bernard S. Lopez.

## Abstract

The DNA damage response (DDR) interrupts cell cycle progression to restore genome integrity. However, unchallenged proliferating cells are continually exposed to endogenous stress, raising the question of a stress-threshold for DDR activation. Here, we identified a stress threshold below which primary human fibroblasts, activate a cell-autonomous response that not activates full DDR and not arrests cell cycle progression,. We characterized this “pre-DDR” response showing that it triggers the production of reactive oxygen species (ROS) by the NADPH oxidases DUOX1 and DUOX2, under the control of NF-κB and PARP1. Then, replication stress-induced ROS (RIR) activates the FOXO1 detoxifying pathway, preventing the nuclear accumulation of the pre-mutagenic 8-oxoGuanine lesion, upon endogenous as well as exogenous pro-oxidant stress. Increasing the replication stress severity above the threshold triggers the canonical DDR, leading to cell cycle progression arrest, but also to RIR suppression. These data reveal that cells adapt their response to stress severity, unveiling a tightly regulated ”pre-DDR” adaptive response that protects genome integrity without arresting cell cycle progression.

## Introduction

Cells are continually challenged by exogenous and endogenous insults that can compromise genome stability, leading to genetic instability and, ultimately, to inflammation, pre-mature ageing and oncogenesis. To counter these stresses, the DNA damage response (DDR) coordinates a network of pathways insuring faithful genome transmission. Defects in DDR result in sensitivity to genotoxic agents, genome instability, neuronal defects, and are frequently associated with cancer predisposition and premature aging (Negrini *et al*, 2010; Gorgoulis *et al*, 2005; Jackson & Bartek, 2009; Kastan & Bartek, 2004; Bartkova *et al*, 2005; Bartek *et al*, 2007; Hoeijmakers, 2009). The DDR is activated at pre/early steps of senescence and tumorigenesis, underlying the role of these processes in preventing cancer initiation (Bartkova *et al*, 2005; Gorgoulis *et al*, 2005; Bartkova *et al*, 2006; Gorgoulis & Halazonetis, 2010; Halazonetis *et al*, 2008).

Following genotoxic stress, the DDR activates cell cycle checkpoints that arrest cell cycle progression and DNA synthesis, giving the DNA repair/recombination machineries the opportunity to repair damaged DNA and the cell to then restart replication on intact DNA matrix. However, in absence of exogenous stress, cells are still routinely submitted to inevitable endogenous stresses, such as replicative stress and oxidative stress, potentially jeopardizing DNA stability. Indeed, replication forks progression is spontaneously hampered by endogenous hindrances (structures difficult to replicate, conflict with transcription, protein tightly bound to DNA, endogenous damages…) (Lambert & Carr, 2013; Carvalho & Lupski, 2016; So *et al*, 2017). In addition reactive oxygen species (ROS), which are spontaneously generated as byproducts of cell metabolism, can directly or indirectly react with DNA (Wallace, 2002), and can also alter replication dynamic (Wilhelm *et al*, 2016; Somyajit *et al*, 2017; Wallace, 2002). Although they are continuously exposed to such chronic low-level endogenous stresses, cells still proliferate and replicate their genome, suggesting that the DDR is not or not fully activated. Thus, this implies that a stress threshold is needed to be reached to fully activate the DDR, as previously proposed (Saintigny *et al*, 2016; Wilhelm *et al*, 2014, 2016), and, as a corollary, that cells either do not respond to low-level stresses, or have developed specific alternative responses.

Cells from patients with DDR syndromes frequently exhibit increased levels of endogenous ROS. In addition, cells deficient in homologous recombination (HR) also exhibit spontaneously high levels of ROS (Wilhelm *et al*, 2016). We hypothesized that such ROS production may participate in an autonomous cellular response to chronic and persistent endogenous genotoxic stress, resulting from repair defects.

Here, we addressed the question of a stress threshold to activate the DDR and of a potential response specific to low stress, below the threshold. We show that cell respond to replicative stress in two distinct phases, adapting the response to stress severity. In primary human fibroblasts, below the DDR activating stress threshold, low replicative stress does not lead to replication inhibition, but induces ROS production driven by the cellular NADPH oxidases DUOX1 and DUOX2, under the control of NF-κB and PARP1. This response protects cells from exogenous exposure to hydrogen peroxide, defining a hormesis-like, low-dose adaptive response. Increasing replication stress (high stress) triggers the canonical DDR (cDDR), which activates the cell cycle checkpoint, arrests DNA synthesis, and also suppresses the replication stress-induced ROS (RIR). Altogether, the cellular response to replication stress can be subdivided in two pathways: An endogenous low-level stress response, licensing DNA synthesis; and a high-stress response that activates the canonical DDR (cDDR) and arrest replication. These data reveal a specific cellular defence response to endogenous/low-level stress, underlining the fine-tuning of the cell response to stress severity.

## Results

### Low replication stress induces ROS

To investigate whether replicative stress actually leads to ROS production, primary human fibroblasts were exposed to hydroxyurea (HU), an inhibitor of ribonucleotide reductase, or aphidicolin (APH), an inhibitor of replicative polymerases. The intracellular ROS level were initially assessed using a 2’,7’-dichlorofluorescein diacetate (DCFDA) fluorescent probe. Both HU and APH treatments led to an increase in intracellular ROS (Figure 1A).

**Figure 1.**
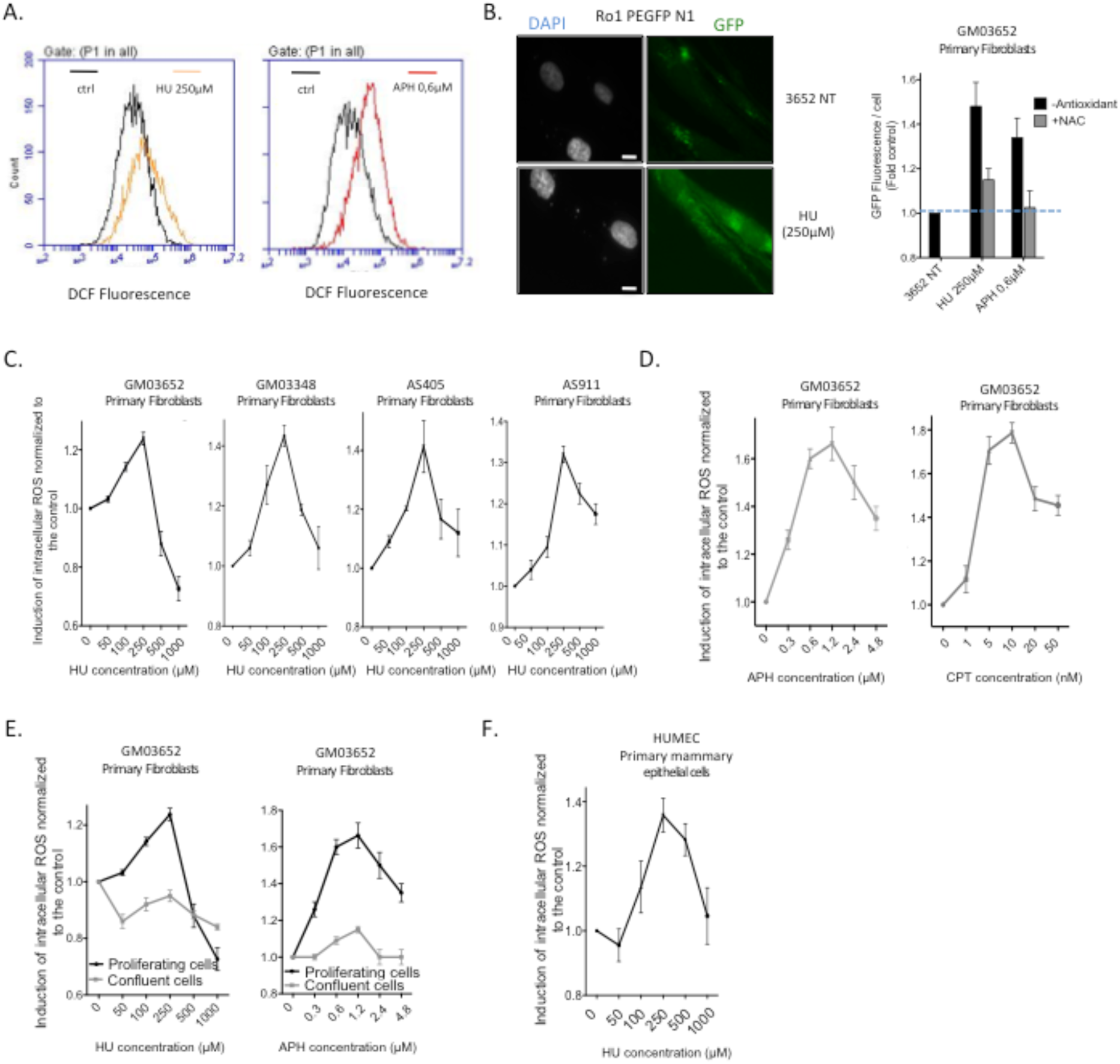
Low replication stress induces cell-regulated ROS production in primary human fibroblasts. **A**. HU- or APH-induced ROS production in primary human fibroblasts was monitored using the DCFDA fluorescent probe and FACS analysis. The shift in the fluorescence peak reveals the induction of intracellular ROS production. **B**. Expression of Ro1 pEGFP-N1 in primary fibroblasts. Left panel: representative images of GFP fluorescence; Scale bars: 10µm. Right panel: quantification of the frequency of ROS positive cells. The histogram represents the means ± SEM (normalized to the control) of four independent experiments, *p < 0.01 compared with the control, as determined by the t-test. **C**. HU dose-dependent induction of ROS production using the DCFDA fluorescent probe in four different primary fibroblast strains. **D**. Impacts of aphidicolin (APH) and camptothecin (CPT), on ROS production in primary fibroblasts. **E**. Cell confluence abrogates HU- and APH-induced ROS production. **F**. The HU-induced ROS production in primary mammary epithelial cells (DCFDA probe).

We confirmed the production of intracellular ROS upon replication stress by a second method, using a plasmid encoding an engineered GFP protein (Ro1 pEGFP-N1) that becomes fluorescent upon oxidation (Dooley *et al*, 2004). Consistent with the data above, exposure of cells to HU increased the frequency of *GFP*^*+*^ cells, an effect that was suppressed by treatment with the antioxidant N-acetylcysteine (NAC) (Figure 1B).

In 4 different human primary fibroblasts strains, replication stress-induced ROS (RIR) levels progressively increased as a function of HU concentration, reaching a peak and then decreased at higher doses (Figure 1 and, Expanded View data Figure S1). Similar dose-response curve shapes were obtained with different replication stress inducers: HU, APH and camptothecin (CPT), a topoisomerase I inhibitor (Figure 1D). Noteworthy, the RIR response was not detectable in confluent non-replicating cells (Figure 1E), suggesting that the production of ROS depends on the proliferation state of the cells. Collectively, these data reveal that ROS are produced in proliferating cells.

Primary epithelial cells also showed peak-shaped production of RIR in response to HU (Figure 1E). Therefore, under each condition and in each cell strain, RIR levels peaked at the same dose (250 µM HU, 0.4 µg/ml APH), suggesting that RIR production constitutes a tightly controlled cell response.

### RIR represents a “pre-DDR” low-stress response

HU doses that induce RIR (≤ 250 µM) did not affect the cell cycle distribution of primary human fibroblasts (Figure 2A). BrdU was efficiently incorporated, indicating cells maintained DNA synthesis at these doses (Figure 2A) while at higher doses (≥ 500 µM HU), where RIR production was not observed (Figs. 1C-F), BrdU incorporation, and thus DNA synthesis, was blocked (Figure 2A). These data suggest that cell cycle checkpoints were activated at higher HU doses, but were not or poorly induced at lower doses that produce RIR. We analysed the activation of DDR markers and found they were significantly induced at higher doses, from 250 to 1000 µM HU (Figure 2B). pCHK1 was poorly induced at low doses (50 and 100 µM HU) and substantially increased at highest doses (Figure 2B). The levels of both phosphorylated Tp53 and p21 protein also substantially increased in primary fibroblasts treated with high HU doses (≥ 250 µM), while the phosphorylation of γH2AX was detected only in primary fibroblasts treated with the highest HU doses (500 and 1000 µM) (Figure 2B). Thus, low HU doses, up to 250 µM, induced RIR production, without fully activating the cDDR and without inducing cell cycle arrest. High HU doses activated the cDDR, leading to concomitant inhibition of DNA synthesis and decrease in RIR levels. The cDDR regulators p53 and ATM have antioxidant functions (Maillet & Pervaiz, 2012; Ditch & Paull, 2012). We therefore tested their impact on RIR levels. While inhibition of p53 or ATM expression using siRNAs did not affect RIR production at 250 µM HU, it abrogated the decrease of RIR at 1 mM HU (Figure 2C).

**Figure 2.**
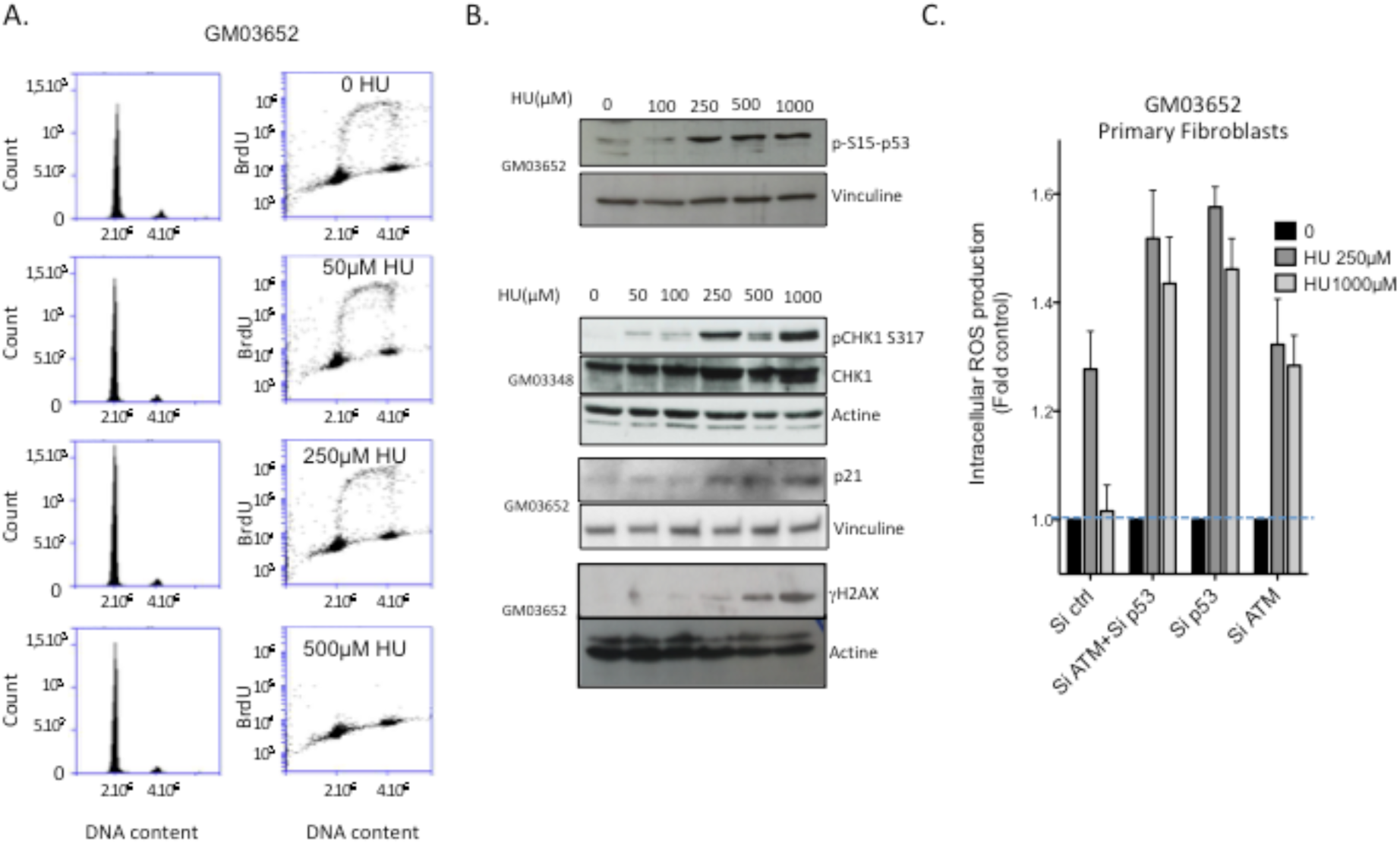
RIR production corresponds to a low-stress response. **A**. Effect of HU on the cell cycle. Primary fibroblasts were treated with increasing concentrations of HU for 72 h, and then PI staining and BrdU pulse labeling were performed to measure cell-cycle distribution (left panel) and DNA synthesis (right panel), respectively. **B**. Immunoblots of phosphorylated p53, CHK1, and γH2AX levels, and p21 in untreated and HU-treated primary fibroblasts after 72 h of treatment. **C**. Impacts of p53 or ATM silencing on the decrease in RIR production induced by 1 mM HU. The data from three independent experiments are presented as the mean ROS production normalized to the control (± SEM).

Thus, replication stress is detected by cells as a function of stress severity. Full cDDR activation, leading to DNA synthesis and cell cycle progression arrests, required a stress threshold equivalent to ≥ 250 µM HU. Above this threshold, DDR markers are strongly activated, and RIR are also suppressed. Below this threshold, certain cDDR markers are not or poorly activated, cells synthesize DNA, they progress through the cell cycle, and RIR production is induced. Therefore, RIR production is part of a specific endogenous/low-level stress response during which cells still replicate their genome. The characterization of the molecular mechanism controlling RIR production becomes thus important.

### RIR are produced by NADPH oxidases

If RIR production is a cell autonomous response, it should be tightly controlled by the cell. NADPH oxidases’ primary function is the cell-regulated production of ROS, which act as secondary messengers (Ameziane-El-Hassani *et al*, 2016; Holmström & Finkel, 2014). Exposure to diphenylene iodonium chloride (DPI), a NADPH oxidase inhibitor (Doussière & Vignais, 1992), abrogated RIR (Figure 3A). Monitoring mRNA expression of the seven identified NADPH oxidases upon exposure of cells to HU revealed that *DUOX1* and *DUOX2* levels increased substantially, in a dose-dependent manner, while *NOX4* and *NOX5* mRNAs were not affected (Figure 3B). *NOX1, NOX2* and *NOX3* mRNAs were not detected in these cells. Of note, cDDR activation (> 250µM HU) does not abrogate the expression of *DUOX1* and *DUOX2*, but instead detoxifies the produced ROS themselves, in a p53/ATM dependent manner (see Figure 2C).

**Figure 3.**
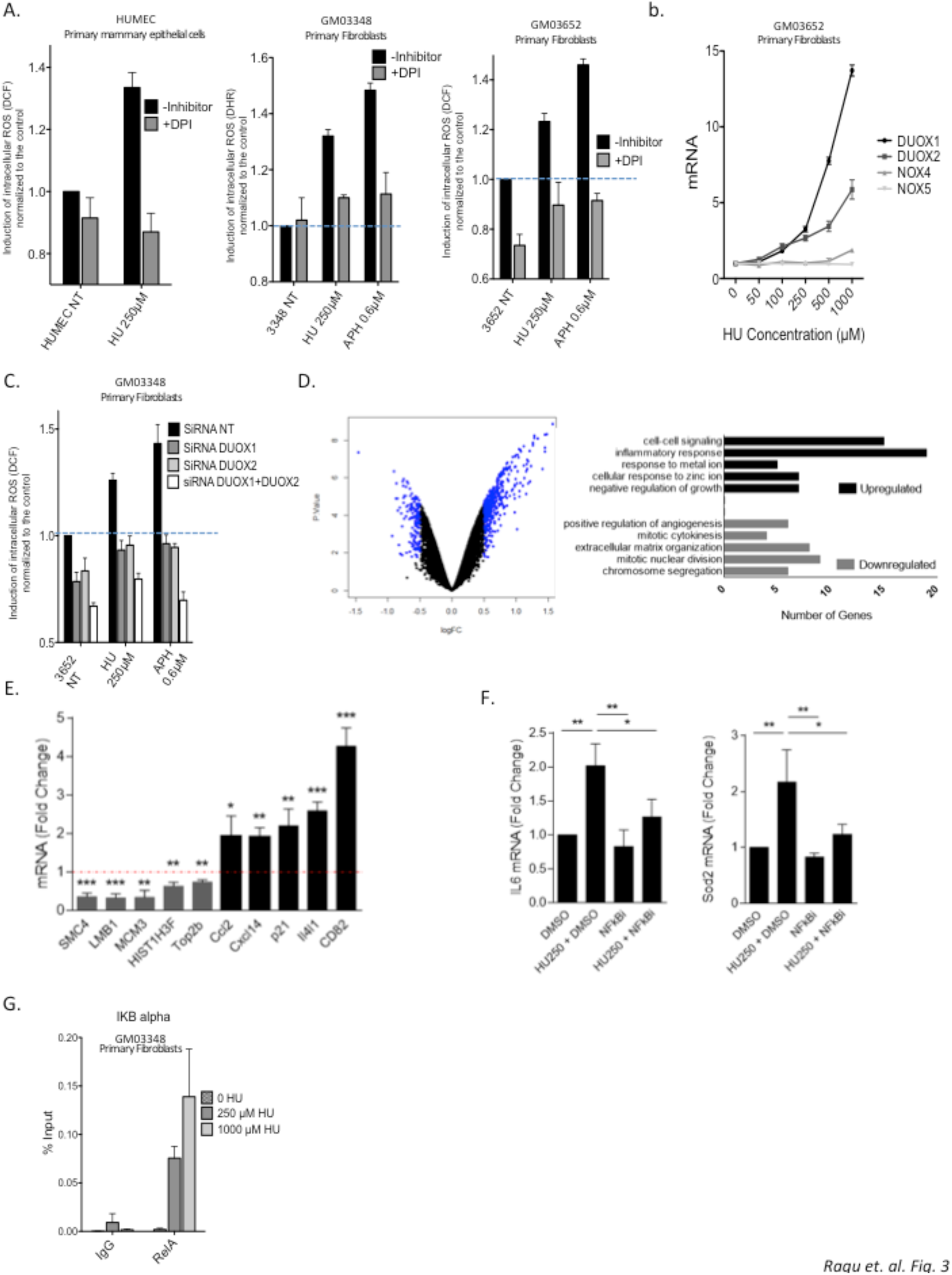
Non-replication blocking doses of HU induce expression of DUOX1 and DUOX2, and NF-κB controlled genes. **A**. Effect of DPI (NADPH oxidase inhibitor) on RIR production, in two different primary fibroblast strains. **b**. Replication stress increases the DUOX1 and DUOX2 mRNA levels in primary fibroblasts. **C**. DUOX1 and DUOX2 silencing impair RIR production in primary fibroblasts. **D**. Transcriptome analysis. Left Panel Vulcano plot from microarray data comparing primary human fibroblasts (GM03348) treated or not with 250 μM HU. The targets with log2 (fold change - FC) > 0.5 and < –0.5, and an adjusted p-value of 0.05 were highlighted as blue points. Right panel: Gene ontology analysis of down- and upregulated genes. **E**. Validation of microarray analysis by real-time RT-PCR analysis for down-regulated (SMC4, LMB1, MCM3, HIST1H3F and Top2b) and up-regulated genes (Ccl2, Cxcl14, p21, Il4l1 and CD82). **F**. Real-time RT-PCR for IL6 and SOD2 using cDNA generated from primary human fibroblasts (GM03348) treated or not with 250 μM HU and DMSO or NF-κB inhibitor (QNZ). **G**. Binding of RelA to an established NF-κB target, the IκB gene promoter.

To determine if *DUOX1* or *DUOX2* are required for RIR production we used siRNA-mediated silencing to knock down (KD) each gene (siRNA efficiency is depicted in Expanded View data Figure S2). KD of either *DUOX1* or *DUOX2* abolished RIR induction (Figure 3C), and simultaneous silencing of both *DUOX1* and *DUOX2* decreased the level of ROS levels even further (Figure 3C).

### Low-dose HU modulates NF-κB-dependent genes

To identify pathways regulating *DUOX1* and *DUOX2* expression upon low replication stress, and therefore RIR, we performed a microarray analysis comparing primary human fibroblasts treated with 250 μM HU with non-treated cells (Figure 3D). Using cutoff values of log2 (fold change - FC) > 0.5 and < –0.5, and an adjusted p-value of 0.05, we identified 152 down- and 416 up-regulated genes in HU-treated cells (Figure 3D and Expanded View data Figure S3A). Gene expression data were then validated by real-time RT-PCR of specifically up- and down-regulated targets (Figure 3E). Gene ontology analysis detected the down-regulation of cell cycle regulating genes, whereas genes involved in inflammation, negative regulation of growth, metabolism of metal and zinc ions, and cell-cell signalling were up-regulated (Figure 3D). 69 genes whose expression was increased by 250 μM HU showed responsive binding sites in their promoter for p65 (RelA), a member of the NF-κB signalling pathway (Expanded View data Figure S3A). By using a selective NF-κB inhibitor, we verified that two classical NF-κB targets (IL6 and SOD2) detected in our microarray analysis were actually up-regulated in a NF-κB-dependent manner after 250 μM HU exposure (Figure 3F).

Chromatin immunoprecipitation (ChIP) using an anti-RelA antibody confirmed that NF-κB was activated following exposure to HU. Indeed, HU stimulated the binding of RelA to the promoter of the *IkB* gene, a target of NF-κB (Figure 3G).

Collectively these data show that 250 µM HU activate the expression of NF-κB dependent genes.

### RIR is induced by NF-κB and PARP1

NF-κB is involved in a variety of physiological and pathological pathways, including cell proliferation and death, immune and inflammatory responses, and tumor immunosurveillance (Hoesel & Schmid, 2013). NF-κB can be induced by genotoxic stresses, including strong replication stress (Wu & Miyamoto, 2008; Christmann & Kaina, 2013). *In silico* analysis revealed a RelA/p65 binding site upstream of the transcription starting sites of both the *DUOX1* and *DUOX2* genes, but none in the other NADPH oxidases encoding genes (http://www.genecards.org/). To test whether NF-κB was involved in RIR production and in up-regulation of *DUOX1* and *DUOX2* expression in primary human fibroblasts exposed to a non-blocking replication stress, we inhibited NF-κB with chemical inhibitors and observed suppression of RIR production induced by either HU or APH (Figure 4A) and repression of stimulation of *DUOX1* and *DUOX2* mRNA expression in HU-treated cells (Figure 4B). RelA accumulates in the nucleus following exposure to 250 µM HU (Figure 4C) and RelA silencing suppressed the HU-increased levels of both the DUOX1 and DUOX2 mRNAs (Figure 4D).

**Figure 4.**
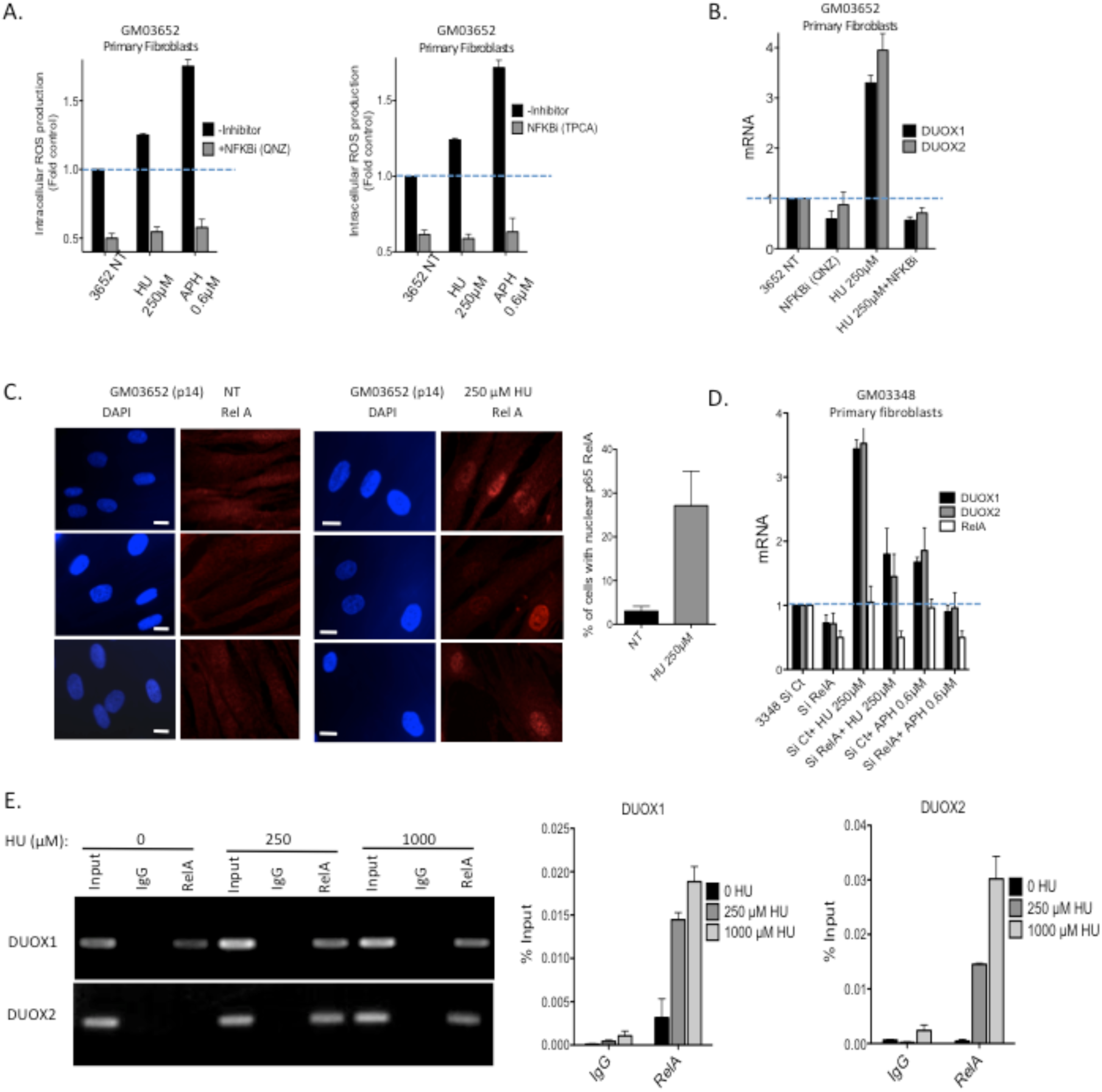
NF-κB controls the production of RIR. **A**. Inhibition of NF-κB with two different inhibitors in primary fibroblasts. **B**. Impact of the inhibition of NF-κB on the expression of DUOX1 and DUOX2 mRNAs. **C**. Immunofluorescence staining for the NF-κB subunit RelA (red) in primary fibroblasts. Left panel: representative photos of immunofluorescence staining for RelA (red) in primary fibroblasts. Nuclei are counterstained with DAPI (blue) Scale bars: 10µm. Right panel: quantification of the nuclear translocation of RelA upon exposure to 250 µM HU. At least 200 cells were counted. **D**. Silencing of RelA inhibits the expression of the DUOX1 and DUOX2 mRNAs. **E**. Binding of RelA to the NF-κB RE sequences located upstream of the TSS in the DUOX1 and DUOX2 genes. Top panel: electrophoresis analysis of the PCR-amplified fragments resulting from the RelA ChIP experiment. IgG: precipitation with a secondary antibody without the primary antibody. RelA: precipitation with the RelA antibody. Low panels: qPCR analysis and quantification, relative to the input. ChIP was performed on primary GM03348 fibroblasts treated with or without HU using an anti-RelA antibody. Primer sequences are listed in the METHODS in Expanded View data. Data from at least three independent experiments are presented (error bars, ± SEM).

ChIP experiments also revealed that RelA binds to NF-κB responsive regions of both *DUOX1* and *DUOX2* promoters, and that this binding was stimulated by exposure of the cells to HU (Figure 4E).

PARP1 is involved in the response to DNA damage, and can activate NF-κB via a mechanism that does not requires PARP1 enzyme activity (Hassa *et al*, 2001; Martín-Oliva *et al*, 2004). PARP1 was therefore a candidate to mediate NF-κB activation in response to DNA stress, leading to the up-regulation of *DUOX1* and *DUOX2* and, *in fine*, to RIR. To test this idea we repressed *PARP1* expression using siRNA KD and observed the loss of induction of RIR by HU and by APH (Figure 5A), while PARP enzyme inhibitors did not affect RIR induction (Expanded View data Figure S4). Silencing PARP1 prevented the nuclear accumulation of RelA (Figure 5B), and the up-regulation of *DUOX1* and *DUOX2*, together with other NF-κB-dependent genes, in response to HU or APH exposure (Figure 5C). Thus, RIR production is controlled by PARP1, which activates the RelA-dependent NF-κB pathway to up-regulate *DUOX1* and *DUOX2*. Then we addressed the question of the role and consequences of RIR induction.

**Figure 5.**
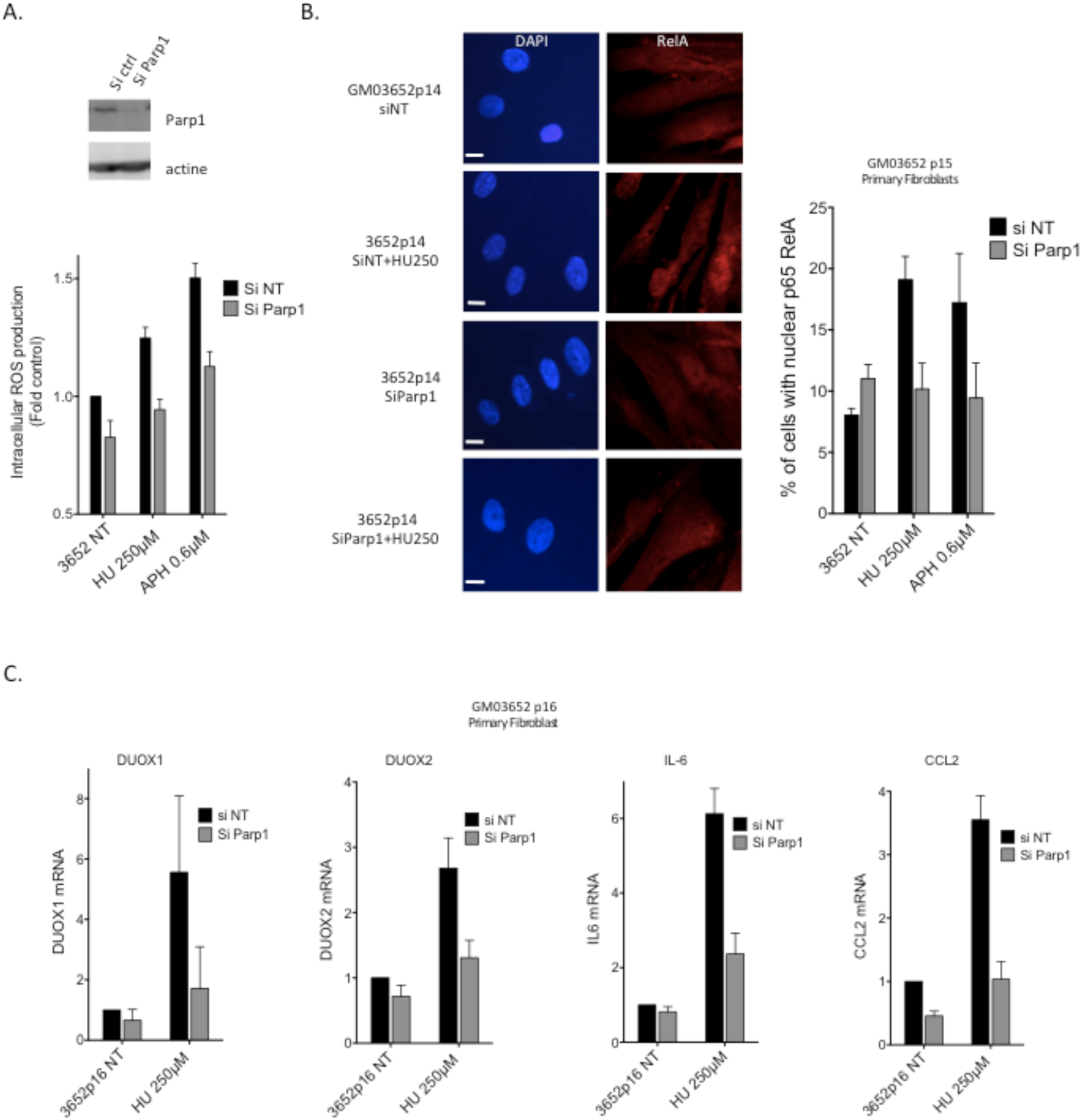
PARP1 controls the production of RIR. **A**. Silencing of PARP1 abolished the induction of HU- or APH-induced RIR. Upper panel: Immunoblot of PARP1 silencing in primary fibroblasts. lower panel: quantification of ROS (DCFA). **B**. Silencing of PARP1 inhibits the HU-induced translocation of RelA in primary fibroblasts. Left panel: representative photos of immunofluorescence staining for RelA (red) in primary fibroblasts. Nuclei are counterstained with DAPI (blue). Scale bars: 10µm. Right panel: quantification of cells with nuclear RelA. Right panel: quantification of the nuclear translocation of RelA. At least 200 cells were counted. The data from four independent experiments are presented. **C**. Impact of PARP1 silencing on the expression (RT-QPCR) DUOX1 and DUOX2 (left panels), and other NF-κB controlled genes (right panels).

### The RIR response protects primary fibroblasts

Given endogenous/low-level stress generates RIR (i.e. ROS), we asked whether it impacts oxidative damage in cells genome. To address this question, we quantified the main pre-mutagenic oxidized base lesion, 8-oxo-Guanine (8-oxoG), in genomic DNA, using isotope dilution high-performance liquid chromatography coupled with electrospray ionization tandem mass spectrometry (HPLC-MS/MS) (Ravanat *et al*, 1998). The pro-oxidant H_2_O_2_ generated high levels of 8-oxoG into genomic DNA (Figure 6A). Low HU doses (50 and 250 µM), which induce ROS (RIR), did not increase, and even decreased the frequency of genomic 8-oxoG (Figure 6B). In contrast, higher HU doses (1000 µM), which do not induce RIR, did not significantly alter the frequency of 8-oxoG into the genomic DNA (compare Figure 6B and 1C-F). These results suggest that the physiological role for RIR is, paradoxically, to protect DNA from the accumulation of pre-mutagenic oxidative DNA alterations.

**Figure 6.**
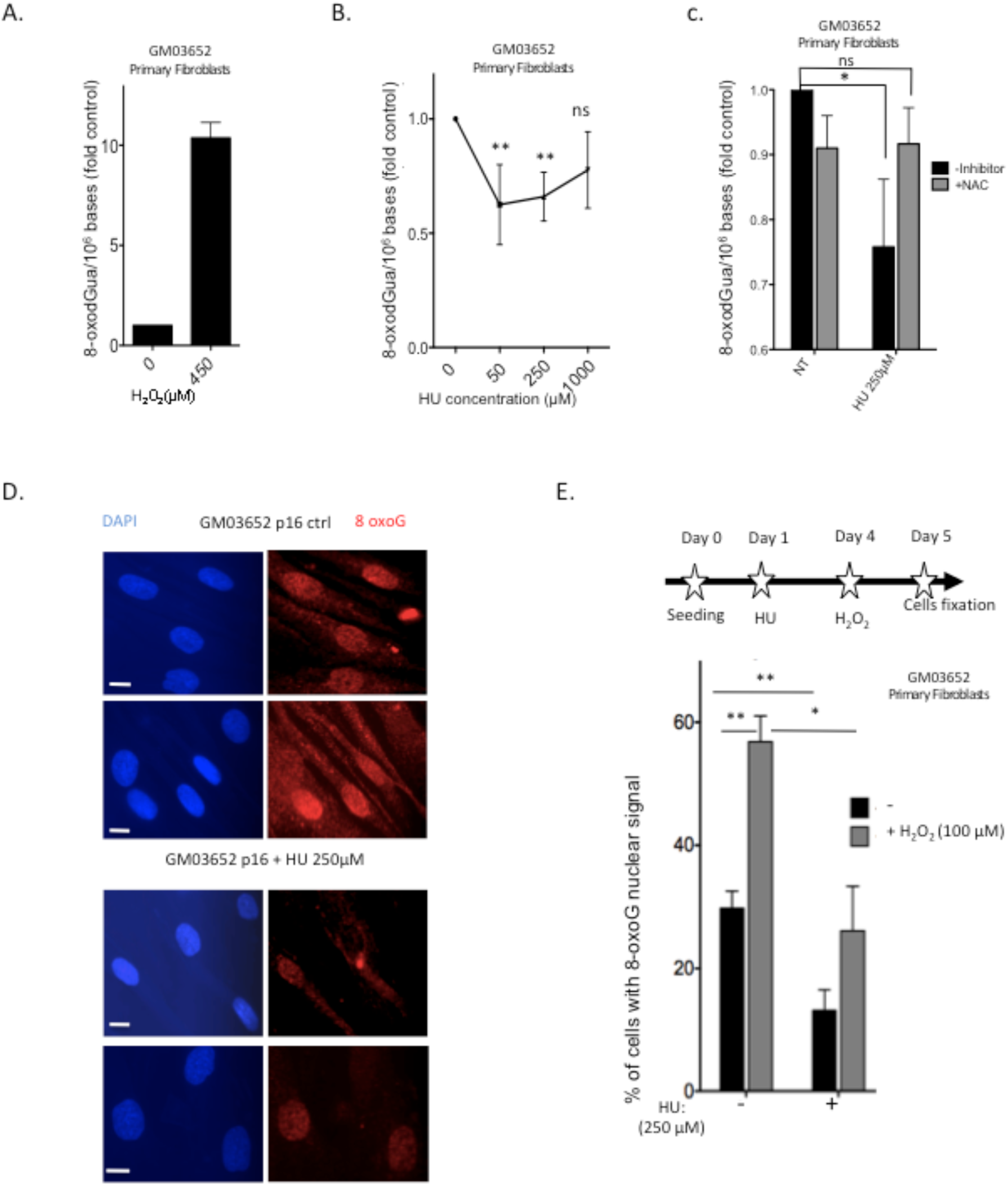
RIR protect primary fibroblasts from endogenous pre-mutagenic oxidative DNA lesions through FOXO1 activation. **A**. Accumulation of 8-oxoG in the genome of primary fibroblasts exposed to 450 μM H_2_O_2_ for 90 min. **B**. 8-oxoG levels in the genome of primary fibroblasts after 72 h exposure to HU. The graphs present 8-oxoG quantification (8-oxoG/million bases). **C**. Effect of NAC on 8-oxoG levels in the genome of primary fibroblasts. **D**. 8-oxoG positive cells (primary fibroblasts) upon 72 h exposure to HU, using an antibody raised against 8-oxoG. Representative photos of immunofluorescence staining for 8-oxoG (red) in primary fibroblasts. Scale bars: 10µm. **E**. Quantification of the frequency of cells with nuclear localization of 8-oxoG, in 250 µM HU-pretreated primary fibroblasts exposed to 100 µM H_2_O_2_. At least 200 cells were counted. Scale bars: 10µm.

To determine whether ROS/RIR protects the genome, we examined the impact of NAC, which abrogates RIR (Figure 1B). We predicted that NAC could have two opposite effects on 8-oxoG accumulation: Based on the above results, the abrogation of RIR could suppress the protection it provides against 8-oxoG accumulation; however, in contrast, a decrease in basal endogenous ROS levels should lead to a decrease in endogenous 8-oxoG frequency in the genome. Unstressed cells exposed to NAC exhibited a slight decrease in 8-oxoG frequency (Figure 6C). However, the substantial decrease in 8-oxoG caused by HU exposure was abolished by NAC treatment, with 8-oxoG levels similar to those in unchallenged control cells (Figure 6C). To confirm these data, we used an anti-8-oxoG antibody and observed that both the immunofluorescence signal intensity of endogenous 8-oxoG (Figure 6D) and the frequency of spontaneous 8-oxoG-positive cells (Figure 6E) decreased upon HU treatment. Exposure to H_2_O_2_ significantly increased the frequency of 8-oxoG-positive cells, but pre-treatment with HU prevented this stimulation (Figure 6E). Thus, RIR induced by pre-treatment with HU prevents the accumulation of endogenous pre-mutagenic 8-oxoG lesions in the genome, including in response to an exogenous pro-oxidant.

To determine the final outcome of on mutagenesis, we performed whole genome sequencing (WGS) of primary human fibroblasts exposed to H_2_O_2_, HU, or both, and compared induced mutations to WGS from untreated cells (the reference sequence) (Figure 7A, 7B and Expanded View data Figure S5). Compared to untreated control genome, H_2_O_2_ generated 398.1 mutations/10^6^ bases, and HU (which lead to replication stress) generated 412.1 mutations/10^6^ bases (Figure 7A), with no hot- or cold-spots regions (Figure 7B). Although mutagenesis induced by the combination of these different treatments was expected to be additive (398.1/10^6^ + 412.1/10^6^ = 810.2 mutations /10^6^ bases), we observed only 346.2 mutations/10^6^ bases (Figure 7A), two fold less than predicted. This observation suggested that HU, while itself mutagenic, protects DNA from over-mutagenicity by H_2_O_2_, consistent with reduced 8-oxoG accumulation (Figure 6B-E). Thus, potential mutagenesis induced by endogenous low-level replication stress is counterbalanced by a mechanism that reduces oxidative stress-induced mutagenesis.

**Figure 7.**
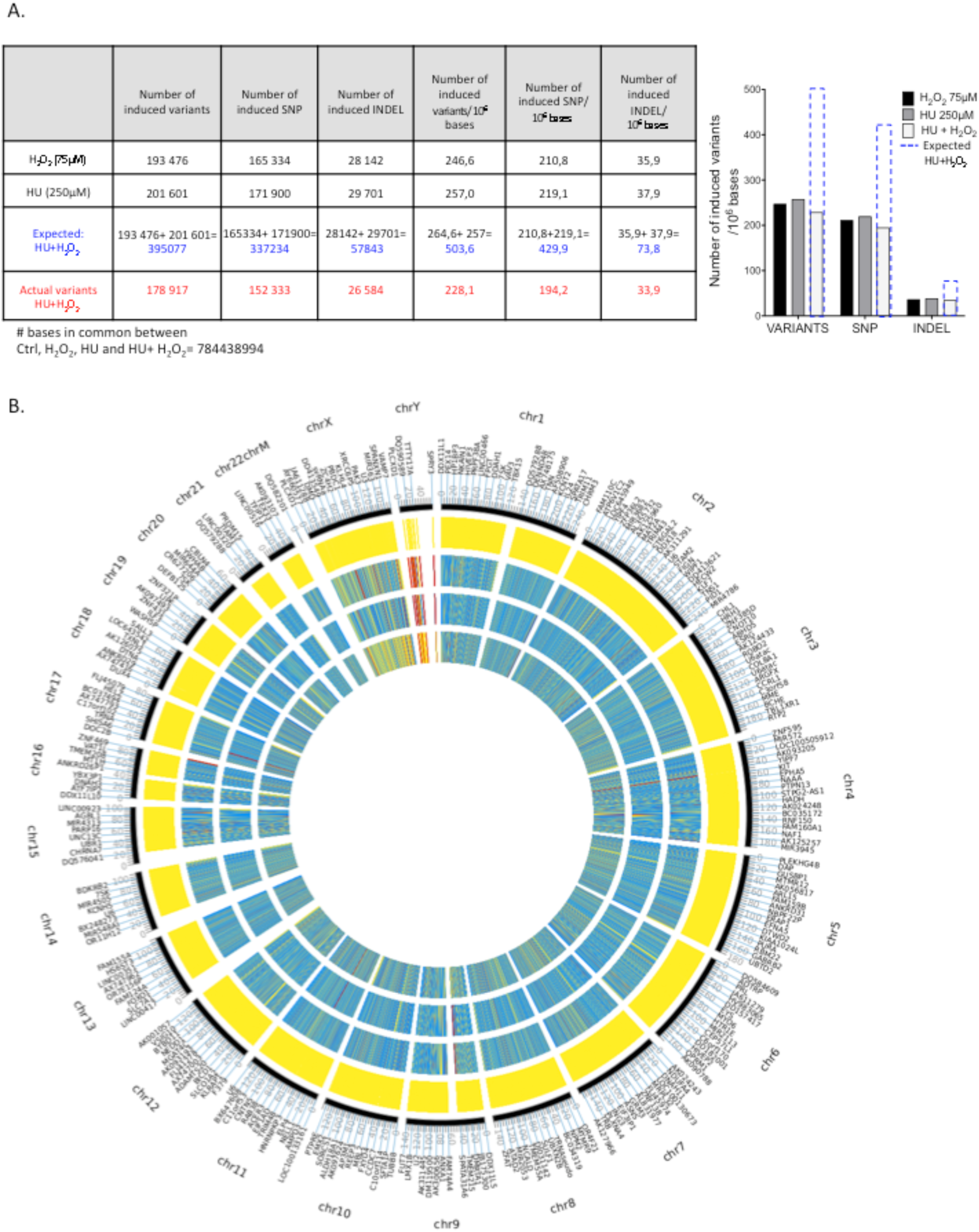
Impact on mutagenesis. **a**. Analysis of the number of actual induced mutations and predicted induced mutations (HU+ H_2_O_2_) compared to untreated controlled cells. Induced mutations: mutations in treated cells subtracted from mutations in the untreated control cells. Left panel: tables with number of mutations in each condition. Right panel: the blue dotted lines represents the predicted mutations resulting from the sum of mutations induced by HU treatment + H_2_O_2_ treatment. Plain bars: actual values. **B**. Mutagenesis analysis by whole genome sequencing of primary fibroblasts exposed to H_2_O_2_, HU or both. To circumvent the sequence coverage heterogeneity, we compared only common covered regions under all conditions (depth 5X). TContro is compared to the reference genome HG19. Genome from treated are compared to the genome of untreated control cells. Yellow: no mutation; blue: heterozygous mutation; red: homozygous mutations. Results are shown from outer to inner side: GM3652 control (no treatment), GM3652 + HU (250µM), GM3652 + H_2_O_2_ (75 µM) GM3652 + HU (250µM) + H_2_O_2_ (75 µM).

### RIR activates the FOXO1 detoxification pathway

Endogenous low-level stress reduces 8-oxoG accumulation in the genome, although, at the same time, it promotes ROS (RIR) production. Two hypotheses could explain this apparent paradox: either RIR induces DNA repair mechanisms, or a ROS detoxification process is activated in a hormesis-type manner, leading to an adaptive response. Immunoblotting of HU-treated cell extracts show that at low doses of HU (50 and 100 µM HU) the level of OGG1, which repairs 8-oxoG, was slightly decreased (Expanded View data Figure S6A), while 8-oxoG was also decreased (Figure 6B). At higher HU doses, OGG1 levels increased (Expanded View data Figure S6A), while the level of 8-oxoG remained unchanged (compare Figure 6B and Expanded View data Figure S6A). The level of MTH1, which removes oxidized nucleotides from nucleotide pools, did not increased but slightly decreased at all HU doses (Expanded View data Figure S6A). *In vitro* repair assays revealed that neither pAPE1 nor OGG1 activities were induced by HU treatment, with OGG1 activity instead being decreased (Expanded View data Figure S6C). Collectively, these data do not support an induction of the DNA repair machinery by HU that could account for the decrease in 8-oxoG in the genome.

Using a candidate approach, we monitored the expression of ROS detoxification genes controlled by the ROS-inducible transcription factors FOXO1. The expression of 4 detoxification genes (*SEPP1, Catalase, GPX1*, and *SOD2*), which all are controlled by FOXO1, was induced by non-replication blocking doses of HU. This induction was suppressed by inhibiting either the NADPH oxidases or NF-κB (Figure 8A). To test if *FOXO1*-controlled ROS detoxification genes are responsible for the reduced levels of 8-oxoG-positive cells induced by a non-replication blocking dose of HU, we used siRNA to KD *FOXO1*. The KD cells showed higher 8-oxoG levels similar to control cells not exposed to HU (Figure 8B). We conclude that that FOXO1-controlled detoxifying genes are part of the endogenous low-level stress response, protecting cells from the accumulation of 8-oxoG in the genome.

**Figure 8.**
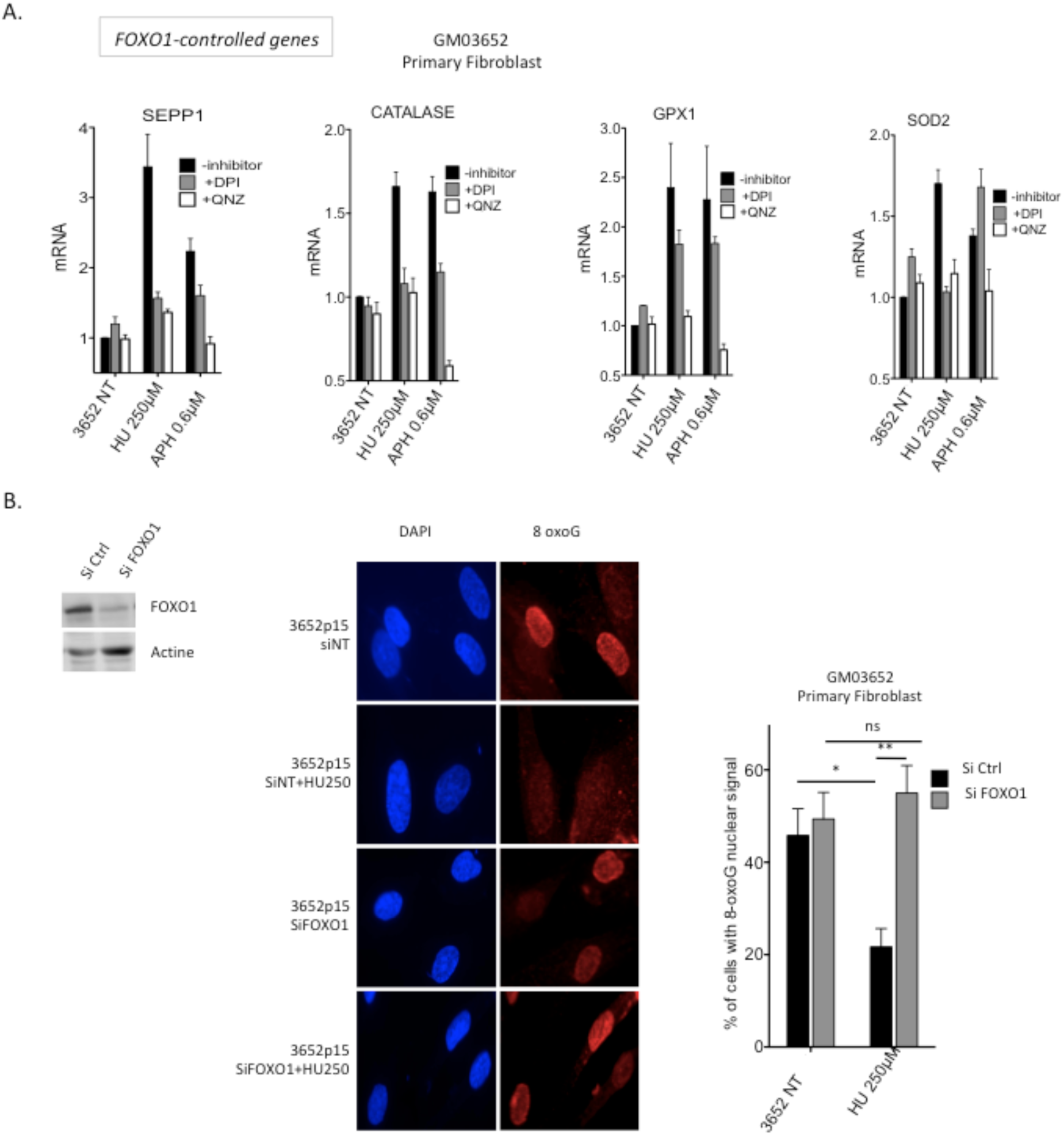
RIR induces the FOXO1 pathway. **A**. Non-replication blocking doses of HU increases the mRNA levels of 4 different detoxification genes (SEPP1, Catalase, GPX1, SOD2) controlled by FOXO1 in primary fibroblasts, and this induction was suppressed by inhibiting the NADPH-oxidases or NF-κB. **B**. Impact of silencing of FOXO1 on the frequency of 8-oxoG positive cells (primary fibroblasts upon) 72 h exposure to HU. Left panel: Immunoblot of FOXO1 silencing in primary fibroblasts. Middle panel: Immunofluorescence staining for the 8-oxoG (red) in primary fibroblasts: representative photos of immunofluorescence staining in primary fibroblasts. Nuclei are counterstained with DAPI (blue). Scale bars: 10µm. Right panel: quantification of the frequency of nuclear 8-oxoG positive cells upon exposure to 250 µM HU after FOXO1 silencing. The data from four independent experiments are presented. At least 200 cells were counted.

## Discussion

Both replication stress and ROS are major endogenous causes of genome instability, senescence and tumorigenesis, and oxidative stress is a significant source of endogenous replication stress (Wilhelm *et al*, 2016; Somyajit *et al*, 2017). Our results reveal that, in a mirror effect, replication stress induces the regulated production of ROS (RIR) that acts as a second messenger in a cell autonomous response. Our data support a bi-phasic model for cell responses to replication stress (Figure 9). At stress levels below a precise threshold (equivalent to 250 µM HU or 0.6 µM APH), the endogenous low-level stress response generates RIR that protect the cell from the accumulation of pre-mutagenic oxidative lesions, such as 8-oxoG. Given the genotoxic potential of ROS, it would appear paradoxical to produce ROS to protect genome integrity. Our results solve this paradox by showing that RIR trigger a ROS detoxification program, resulting in an adaptive response in human primary cells.

**Figure 9.**
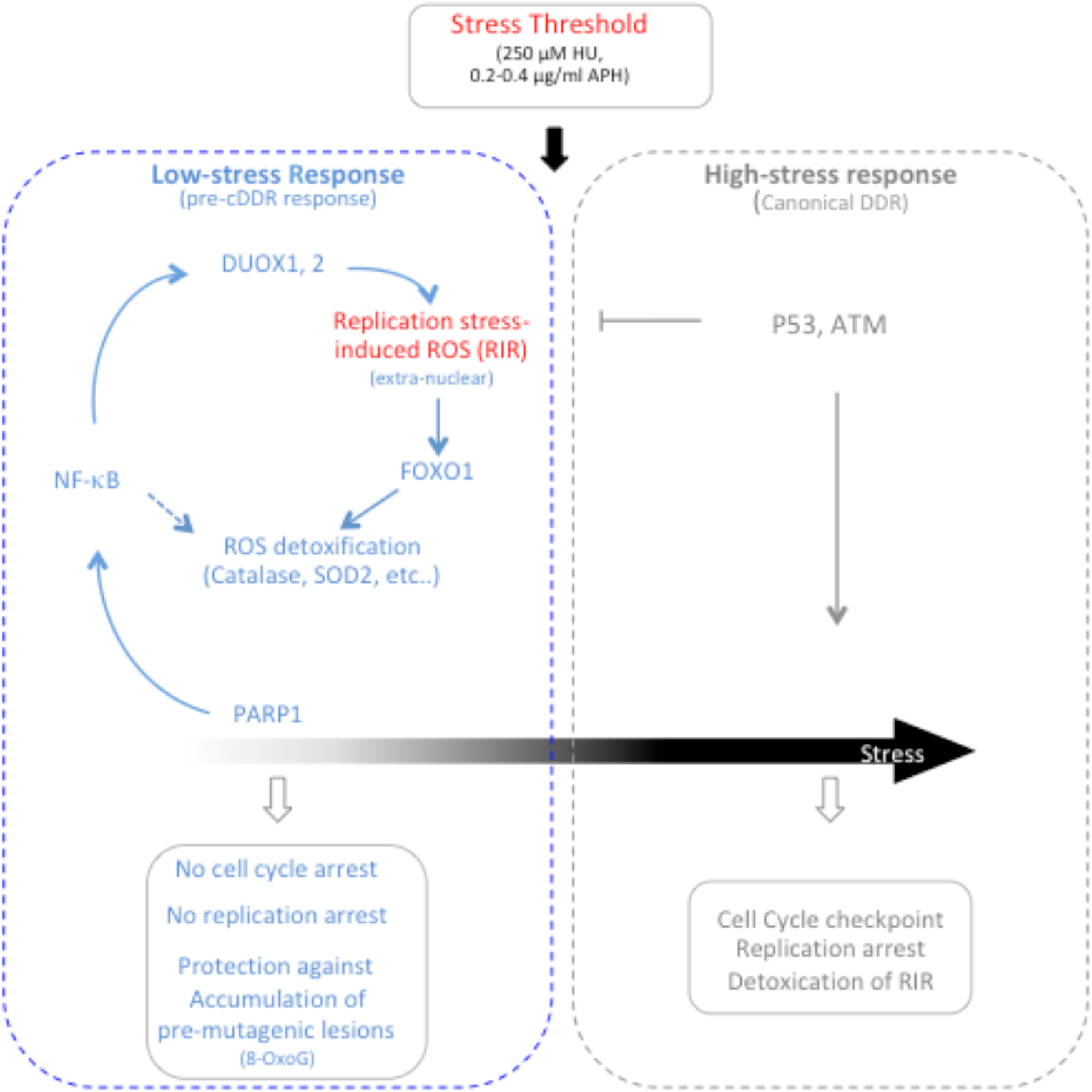
Primary cells adapt their response to replication stress intensity through the low stress response and the high response: A threshold of stress intensity must be reached to activate the high-stress response, which corresponds to the cDDR, and leads to DNA synthesis inhibition and cell cycle arrest. Below this stress intensity threshold, cells engage the endogenous low-level stress response, which does not repress DNA synthesis and cell progression. The endogenous low-level stress response regulates the production of ROS (RIR), protecting the genome from the accumulation of pre-mutagenic lesions, such as 8-oxoG. The endogenous low-level stress response may be considered a “pre-cDDR” response that allows cells to proliferate, thus replication their DNA, but generates RIR that induces the FOXO1 detoxifying program. RIR acts by triggering a bi-phasic, hormesis-like detoxification response. The cell itself regulates extranuclear RIR production through the PARP1/NF-κB pathway, which controls the expression of DUOX1 and DUOX2. When the stress intensity reaches a specific threshold, the endogenous low-level stress response is repressed and replaced by the cDDR, which arrests DNA synthesis, and results in p53 and ATM-dependent detoxification of RIR.

Because ROS represent a threat to the cell, RIR should be tightly controlled. Here we demonstrate that cell control RIR production through the PARP1/ NF-κB pathway(s) that regulates the expression of *DUOX1* and *DUOX2* genes. Because of the potential risks of ROS, increasing ROS production with stress severity would likely *in fine* jeopardize DNA and other cellular components. Therefore, when stress intensity reaches a threshold, the endogenous low-level stress response is replaced by the cDDR, which arrests DNA synthesis and results in p53-and ATM-dependent detoxification of RIR. Hence, the endogenous low-level stress response could be considered as a “pre-cDDR” response, which allows cells to replicate their DNA, but induces the FOXO1 detoxifying program through the cell-controlled production of RIR, protecting genome integrity.

We propose that the Low-stress response helps cells cope with endogenous/low-level replication stress. Some oxidized bases can block DNA polymerase progression (Wallace, 2002) and endogenous ROS can also alter replication dynamics through oxidization of proteins such as the ribonucleotide reductase that synthetize nucleotides (Wilhelm *et al*, 2016). Therefore, RIR detoxification should counterbalance low replication stress by decreasing endogenous other causes of replication stress. Increasing the stress level would increase DNA damage frequency and exhaust nucleotide pools, at which point replication arrest would be triggered by cDDR. In this latter condition, RIR become superfluous and potentially dangerous, and it is inactivated by ATM and p53.

HU treatment is used on patients suffering from myeloproliferative syndromes, to limit cell proliferation. For instance, chronic myelomonocytic leukemia, a severe clonal hematopoietic malignancy, is the most frequent myelodysplastic syndrome/myeloproliferative neoplasm. Most patients receive symptom-adapted treatments, such as HU in the most proliferative forms of the disease (Solary & Itzykson, 2017). Therefore, determining whether these treatments actually activate or not the NF-κB/FOXO1-protective “low-stress” pathway(s) represents thus a future crucial issue, to evaluate the consequences of the chronic exposure to HU of the patients.

In worms, NADPH oxidases trigger redox signaling that favors resistance to oxidative stress and promotes longevity (Ewald *et al*, 2017), supporting our interpretation. In humans, while various NADPH oxidases are often upregulated in cancers, only DUOX1 and DUOX2, which produce RIR in primary cells (see Figure 3A-C), are lost in different kinds of tumors (Little *et al*, 2017), highlighting the protective effects of the DUOX1- and DUOX2-controlled ROS. In line with this, antioxidants (including NAC that abrogates RIR), which have been proposed to protect against carcinogenesis, in fact foster lung carcinomas and metastasis (Breau *et al*, 2019; Le Gal *et al*, 2015; Wiel *et al*, 2019). More generally, defect in each of the players of the “Low-stress” response described here (NF-κB, PARP1, FOXO1, *DUOX1* and *DUOX2*) share common phenotypes, such as defects in cell homeostasis and metabolism, ageing and cancer predisposition (Ewald, 2018; Tia *et al*, 2018; Tilstra *et al*, 2011; Vida *et al*, 2016; Little *et al*, 2017; Park & Hong, 2016; Pires *et al*, 2018).

The data presented here reveal a specific cellular response to low/endogenous stress, different from the canonical DDR, thus underlining the capacity of cells to adapt their responses to the severity of the replication stress they experience. Cells are chronically exposed to unavoidable endogenous low-level stresses throughout their lifespan, in contrast to acute exposure to severe stress. The pathway we have identified and characterized should thus play an essential role in the maintenance of genome stability in the face of such ineluctable low/endogenous-level stress.

## Supporting information

supplementary daata

## Material and Methods

### Cell culture and treatments

Cells were grown at 37°C with 5% CO_2_ in modified Eagle’s medium (MEM). Primary human fibroblasts were grown in MEM (Gibco, Life Technologies) supplemented with 20% fetal calf serum (FCS; Lonza Group, Ltd.). Primary human mammary epithelial cells (HuMECs) were provided by Thermo Fisher Scientific and cultured according to the manufacturer’s recommendations. Primary fibroblasts were exposed to HU, APH or CPT for 72 h at 37°C. For the antioxidant treatment, primary fibroblasts were exposed to 2 mM NAC (Sigma-Aldrich, St. Louis, MO, USA) for 72 h. For the inhibition of NF-κB, fibroblasts were treated with 1 µM QNZ (EVP4593) or 20 µM TPCA-1 (Santa Cruz and Selleckchem, respectively) for 72 h. Primary fibroblasts were transfected with a DUOX1-targeted siRNA (siDUOX1, si On-Target Plus SMARTpool L-008126-00) purchased from Dharmacon Inc. (Lafayette, CO), a DUOX2-targeted siRNA, a RelA-targeted siRNA, or a control non targeting siRNA purchased from Santa Cruz Biotechnology, or a FOXO1-targeted siRNA (GAGCGUGCCCUACUUCAAGGA) using the Amaxa(tm) Basic Nucleofector(tm) Kit for Primary Mammalian Fibroblasts (Lonza), according to the manufacturer’s protocol. For the inhibition of p53, ATM or PARP1 primary fibroblasts were transfected with a p53-targeted siRNA (siTP53, si On-Target Plus SMARTpool L-003329-00), ATM-targeted siRNA (siATM, si On-Target Plus SMARTpool L-003201-00), control nontargeting siRNA (si On-target Plus nontargeting pool D-001810-10-05) all of which were purchased from Dharmacon Inc. (Lafayette, CO), or PARP1-targeted siRNA (siPARP1, Santa Cruz Biotechnology) using the Interferin transfection reagent (Invitrogen), according to the manufacturer’s protocol.

### Measurement of cellular ROS production by FACS

Cellular ROS production was measured using a CM-H2DCFDA (2’,7’-dichlorofluorescein diacetate) (Life Technologies, USA) assay kit, according to the manufacturer’s protocol. Approximately 10^5^ cells/well were plated in 6-well plates and incubated at 37°C with 5% CO_2_. After 2 (cell lines) or 3 days (primary fibroblasts and primary mammary epithelial cells), the cells were rinsed with PBS and incubated with 10 µM CM-H2DCFDA in DMEM supplemented with 1% FBS for 45 min at 37°C in the dark. The cells were trypsinized and resuspended in DMEM supplemented with 1% FBS. The pelleted cells were washed again, and the live pelleted cells were resuspended in PBS and analyzed with a BD Accuri C6 flow cytometer (BD Biosciences, San Diego, CA) equipped with an FL1 laser (515–545 nm). The data are presented as the mean percentages of four independent experiments.

### Measurement of cellular ROS production using the green fluorescent protein RoGFP

RoGFP was expressed in primary fibroblasts using modified pEGFP-N1 as the expression vector and JetPei as the transfection reagent. After cells were incubated in culture medium treated with or without HU or APH for 72 h at 37°C, the cells were washed twice with Hanks’ balanced salt solution. The cells were incubated with DAPI (1 μg/ml) and imaged using a microscope. The images were captured using the 63x oil immersion objective of a motorized Axio Imager.Z2 epifluorescence microscope (Carl Zeiss) equipped with a high-sensitivity cooled interline CCD camera (Cool SNAP HQ2; Roper Scientific) and a PIEZO stage (Physik Instrumente). Images were acquired using MetaMorph software (Molecular Devices). In each case, 300–500 cells were analyzed per condition.

### Western blot analysis

Cells were suspended in lysis buffer (8 M urea, 1 M thiourea, 4.8% CHAPS, 50 mM DTT, 24 mM spermine dehydrate, a protease inhibitor cocktail (Complete Lysis Buffer; Roche, Meylan, France), and 0.1 mM Na_3_VO_4_) and proteins were extracted after repeated mechanical disruption of the lysate through a needle attached to a 0.3 ml syringe. After the samples were incubated for 1 h at room temperature, they were cleared by centrifugation at 13000 rpm. For each blot, equal amounts (30 mg) of protein from each sample were loaded on the gel. Electrophoretic separation, transfer to a nitrocellulose membrane and antibody probing were performed using standard techniques. The proteins were visualized using the ECL Western Blotting System. Actin was probed with a 1:1000 dilution of a specific antibody (Sigma-Aldrich), and the non-phosphorylated and phosphorylated forms of Chk1 were detected using 1:500 dilutions of anti-P (S317)-Chk1 antibodies (Cell Signaling Technology). A 1:500 dilution of the anti-phospho-histone H2A.X (Ser139) antibody (Merck Millipore) was used to detect phosphorylated histone H2A.X levels and phosphorylated p53 was detected using anti-P (S15)-p53 antibodies (Abcam). OGG1 was probed with a specific antibody at 1:1000 (Novus Biologicals). MTH1 was probed with 1:500 dilutions of a specific antibody (Invitrogen). FOXO1 was probed with 1:500 dilutions of a specific antibody (Cell Signaling) and PARP1 or p21were detected using 1/500 dilutions of specific antibody (Santa Cruz Biotechnology).

### 8-oxoG measurement

Genomic DNA was extracted and enzymatically digested using an optimized protocol that minimizes DNA oxidation during the procedure(Ravanat *et al*, 2002). Then, 8-oxoG levels were quantified using isotope dilution high-performance liquid chromatography coupled with electrospray ionization tandem mass spectrometry (HPLC-MS/MS) as previously described(Ravanat *et al*, 1998); ^15^N_5_-8-oxoG was used as the internal standard. In addition to the mass spectrometric detector, the system was also equipped with a UV detector that was set at 260 nm to measure the quantity of normal nucleosides. The results are expressed as the number of 8-oxoG per million normal nucleosides.

### Immunofluorescence

The cells were grown on glass coverslips, fixed with 2% paraformaldehyde and permeabilized with 0.5% Triton X-100. After blocking with 3% BSA in PBS containing 0.05% TWEEN 20, the cells were incubated with anti-RelA (sc-372, Santa Cruz Biotechnology) primary antibodies diluted in PBS containing 3% BSA and 0.05% TWEEN 20. After washing with PBS containing 0.05% TWEEN 20, the cells were incubated with Alexa Fluor 568-conjugated anti-mouse secondary antibodies (Invitrogen, Molecular Probes) and stained with DAPI. <u>For 8-oxoG detection in nuclear</u> <u>DNA</u>, after permeabilisation with 0,5% Triton X-100, cells were denatured by 2N HCl to allow access of the antibody to the chromatin. Then, cells were washed three times in PBS and neutralized with 50 mM Tris–HCl pH 8.8 before proceeding to the blocking with 2% fetal calf serum in PBS containing 0.05% TWEEN 20. The cells were incubated with mouse anti-8-oxo-dG (clone N45.1, 1:100, ab48508 Abcam). Secondary antibodies, goat anti-mouse-IgG Alexa 568, (Invitrogen) were used. Nuclear DNA was counterstained with DAKO DAPI. Images were captured using a Zeiss motorized Axio Imager.Z2 epifluorescence microscope with a 63x/1.4 NA oil immersion objective equipped with a Hamamatsu camera. Data were acquired using AxioVision (4.7.2.).

### Cell cycle analysis and BrdU incorporation

Cells were incubated in culture medium treated with or without HU for 72 h at 37°C, and 5-bromo-2-deoxyuridine (BrdU, Sigma) was added to the culture media at a final concentration of 10 μM for 20 min. Pelleted cells were detached with trypsin, fixed with 80% ethanol, and resuspended in 30 mM HCl/0.5 mg/ml pepsin. BrdU was immunofluorescently labeled with a mouse anti-BrdU antibody (DAKO, clone Bu20a) and a fluorescein-conjugated donkey anti-mouse antibody (Life Technologies), and the cells were stained with propidium iodide (PI; 25 μg/ml) in the presence of ribonuclease A (50 μg/ml). Flow cytometry analyses were performed using an Accuri C6 flow cytometer (BD Biosciences).

### RNA extraction and quantitative RT-PCR (TaqMan)

Total RNA was isolated using the Macherel-Nagel NucleoSpin RNA kit, according to the manufacturer’s instructions. The cDNAs were generated from 2 μg of total RNA using random hexamers and RevertAid Premium Reverse Transcriptase (Thermo Fisher Scientific). The following primers were used for TaqMan® probe-based assay (Applied Biosystems) was used for qPCR of DUOX1 (Hs00213694_m1), DUOX2 (Hs00204187_m1), NOX4 (Hs00276431_m1), NOX5 (Hs00225846_m1), RELA (Hs00153294_m1), SEPP1 (Hs01032845_m1), CATALASE (Hs00156308_m1), GPX1 (Hs00829989_gH), SOD2 (Hs00167309_m1), TXNRD1 (Hs00917067_m1), NQO1 (Hs01045993_g1), FTL (Hs00830226_gH), MGST1 (Hs00220393_m1) and BETA-ACTIN (Hs99999903_m1) served as the internal control. The sequences of the primers used for SYBR assays are in Expanded View l data Table S7. Quantitative RT-PCR was performed using the Applied Biosystems 7300 Real-Time PCR System. All experiments were performed in triplicate.

### ChIP and quantitative PCR (ChIP-qPCR)

Primary fibroblasts were treated with or without 250 μM or 1000 μM HU, cross-linked with 1% formaldehyde 10 min, and then incubated with 125 mM glycine for 5 min. Cells were washed with ice-cold PBS, collected by scraping and centrifuged at 1000 x g for 5 min at 4°C. The supernatant was removed and the pellets were resuspended in lysis buffer (50 mM Hepes pH 8.0, 140 mM NaCl, 1 mM EDTA, 1% Triton X-100, 0.1% sodium deoxycholate, 0.5% SDS, and freshly added protease inhibitors) and incubated on ice for 10 min before sonication. Sonicated chromatin was diluted in radioimmunoprecipitation assay (RIPA) buffer (50 mM Tris, pH 8.0, 1% Triton X-100, 1 mM EDTA, 150 mM NaCl, 0.1% sodium deoxycholate, and 0.05% SDS, freshly added protease inhibitors) and incubated with pretreated beads and 5 μg of the RelA antibody (NF-κB p65 C-20; Santa Cruz Biotechnology) overnight at 4°C. Beads were then washed with washing buffer 1 (20 mM Tris-HCl, pH 8.0, 1% Triton X-100, 2 mM EDTA, 150 mM NaCl, 0.1% SDS, and freshly added protease inhibitors) for 5 min, followed by sequential washes with buffer 2 (20 mM Tris-HCl, pH 8.0, 1% Triton X-100, 2 mM EDTA, 300 mM NaCl, 0.1% SDS, and freshly added protease inhibitors), buffer 3 (10 mM Tris-HCl, pH 8.0, 250 mM LiCl, 1% NP40, 1 mM EDTA, 1% sodium deoxycholate, and freshly added protease inhibitors) and TE buffer (10 mM Tris-HCl, pH 8.0, 1 mM EDTA; two times) for 5 minutes each. One hundred microliters of elution buffer (10 mM Tris-HCl, pH 8.0, and 1 mM EDTA) were added and the beads were incubated with RNase A (400 μg/ml) and NaCl (600 mM) in a Thermo mixer for 1 h at 37°C at 1400 rpm. Then, proteinase K (400 μg/ml) and 1% SDS were added and the mixture was incubated at 65°C overnight with agitation. Beads were precipitated and the supernatants were treated with phenol/chloroform/isoamyl alcohol followed by centrifugation at 13000 x g for 5 min. The supernatants (aqueous phase) were incubated with 300 mM sodium acetate and cold ethanol for 1 h at -80°C, followed by centrifugation at 13000 x g for 30 min. The DNA pellets were washed with 70% ethanol and resuspended in nuclease-free water. The DNA was subjected to qPCR identification with PowerUp SYBR Green Master mix (Applied Biosystems) using an Applied Biosystems 7300 Real-Time PCR System. The data were analyzed with a standard curve-based method. Reference qPCR primers:

Human DUOX1: GPH1004464(-)02A (Qiagen)

Human DUOX2: GPH1018346(-)08A (Qiagen)

### Whole genome analysis

200 ng of genomic DNA was sheared with the Covaris E220 system (LGC Genomics / Kbioscience). DNA fragments were end-repaired, extended with an ‘A’ base on the 3′end, ligated with indexed paired-end adaptors (NEXTflex, Bioo Scientific) using the Bravo Platform (Agilent), amplified by PCR for 6 cycles and purified with AMPure XP beads (Beckman Coulter). The final libraries were pooled and sequenced using the onboard cluster method, as paired-end sequencing on S4 flowcell (2×150 bp reads) on Illumina NovaSeq-6000 sequencer at GR (Illumina, San Diego, CA).

In order to compare them, we focused the analysis of small variant mutations in genome regions that were commonly covered under all conditions. qRaw reads were mapped against hg19 reference sequence with only chromosomes 1, 2, 3, 4, 5, 6, 7, 8, 9, 10, 11, 12, 13, 14, 15, 16, 17, 18, 19, 20, 21, 22, X, Y, MT (NC_012920.1. All haplotypes, chrUn and chr_gl sequences were removed, in order to avoid multiple mappings. Reads alignment was performed using bwa-0.7.12 (mem) and converted to bam with samtools-1-1. Then, Picard-1.121 (SortSam) was used to sort bam by coordinates and duplicate fragments were marked using Picard-1.121 (MarkDuplicates). Next, bam files were merged together with Picard-1.121 (MergeSamFile) and indexed with samtools-1-1. Initial alignments were refined by local realignment using GATK-3.5 (RealignerTargetCreator, IndelRealigner). To finish, a base recalibration was applied on bam files with GATK-3.5 (BaseRecalibrator, PrintReads). The SNPs and INDELs were called with GATK-3.5 (HaplotypeCaller).

In order to compare the variants across the samples, only the regions covered with at least 5X in all samples were kept (bedtools 2.26 - intersectbed). To finish, the variants detected in the control sample were removed. The variant comparisons were made with a in house script in R (v3.5.1) and the package VennDiagram. The Circos plot was made with CircosVCF.

### Microarray Assay

Gene expression analysis were performed with Agilent® SurePrint G3 Human GE 8×60K Microarray (Agilent Technologies, AMADID 39494) with the following dual-color design: the test samples were labeled with Cy5 whereas the control samples were labeled in Cy3 using the two-color Agilent labeling kit (Low Input Quick Amp Labeling Kit 5190-2306) adapted for small amount of total RNA (100 ng total RNA per reaction). Hybridization were then performed on microarray using 825 ng of each linearly amplified cRNA labeled Cy3 or Cy5 sample, following the manufacturer protocol (Agilent SureHyb Chamber; 1650ng of labeled extract; duration of hybridization of 17 hours; 40 µL per array; Temperature of 65 °C). After washing in acetonitrile, slides were scanned by using an Agilent G2565 C DNA microarray scanner with defaults parameters (100° PMT, 3 µm resolution, at 20°C in free ozone concentration environment. Microarray images were analyzed by using Feature Extraction software version (10.7.3.1) from Agilent technologies. Defaults settings were used.

### Microarray data processing and analysis

Raw data files from Feature Extraction were imported into R with LIMMA (Smyth, 2004, Statistical applications in Genetics and molecular biology, vol3, N°1, article3), an R package from the Bioconductor project, and processed as follow: gMedianSignal and rMedianSignal data were imported, controls probes were systematically removed, and flagged probes (gIsSaturated, gIsFeatpopnOL, gIsFeatNonUnifOL, rIsSaturated, rIsFeatpopnOL, rIsFeatNonUnifOL) were set to NA. Intra-array normalization was performed by loess normalization, followed by a quantile normalization of both Cy3 and Cy5 channel. Then inter-array normalization was performed by quantile normalization on M values. To get a single value for each transcript, taking the mean of each replicated probes summarized data. Missing values were inferred using KNN algorithm from the package ‘impute’ from R bioconductor. Normalized data were then analyzed. To assess differentially expressed genes between two groups, we start by fitting a linear model to the data. Then we used an empirical Bayes method to moderate the standard errors of the estimated log-fold changes. The top-ranked genes were selected with the following criteria: an absolute fold-change > 2 and an adjusted p-value (FDR) < 0.05. Microarray Gene Ontology analysis and transcription factors binding sites were performed using the DAVID platform (Huang *et al*, 2009a, 2009b).

### Statistical analyses

Student’s t-test was used to compare differences between two groups. One-way ANOVA was used for comparisons among three or more groups. P < 0.05 was considered statistically significant.

## Accession numbers

The transcriptome microarrays data have been uploaded to ArrayExpress and the accession number is E-MTAB-8605.

The WGS data have been uploaded to SRA and the accession number is PRJNA595802.

## Acknowledgments

We thank Corinne Dupuy and Stéphane Koundrioukoff for helpful discussions and advice on NADPH-oxidases. We thank Roger Tsien (San Diego, USA) for the generous gift of the RoGFP-based plasmids. We thank Noémie Pata-Merci and Guillaume Meurice (AMMICa; INSERM UMS23, CNRS US3655) for transcriptome primary analysis. Whole exomedata analysis was performed by GenoSplice technology (www.genosplice.com). This work is supported by grants from Ligue Nationale contre le cancer “Equipe labellisée 2020” (BSL and ES), ANR (ANR-16-CE12-0011-02, ANR-16-CE18-0012-02), AFM-Téléthon and INCa (Institut National du Cancer 2018-1-PLBIO-07 for BSL and PRT-K16 057 for ES). French National Research Agency, ANR-18-CE44-0008 to A.A.I., CIENCIACTIVA/CONCYTEC Doctoral Fellowship to G.Z. Whole exomedata analysis was performed by GenoSplice technology (www.genosplice.com) The authors declare having no competing financial interests. The authors declare having no conflict of interests.

## Author contributions

SR, GMR, AB, CG, SC and ED performed the experiments. SC and J-L R performed the 8-oxoG dosages. GZ and AAI performed APE1 activity assay. All authors interpreted and discussed the experiments and results. SR and BSL wrote the main manuscript, and all authors corrected it. SR and BSL prepared the figures and Expanded View data.

