## supplementary daata for "An adaptive pre-DNA-damage-response protects genome integrity"

### Expanded View data.

**S1. Individual experiments showing HU-induced ROS.**

**S2. Efficiency of silencing of DUOX1 and DUOX2 mRNA in primary fibroblasts.**

**S3. Microarray analysis comparing primary human fibroblasts treated with HU and untreated primary fibroblasts cells.**

**S4. Impact of PARP inhibitors on RIR production.**

**S5. Whole genome sequencing data.**

**S6. MTH1 and OGG1 proteins AND OGG1 and APE1 activity upon HU exposure.**

**S7**: List of primers used for SYBR realtime RT-PCR.

Supplementary data S1.

**GM03652**

Figure S1. Individual experiments showing HU-induced ROS.

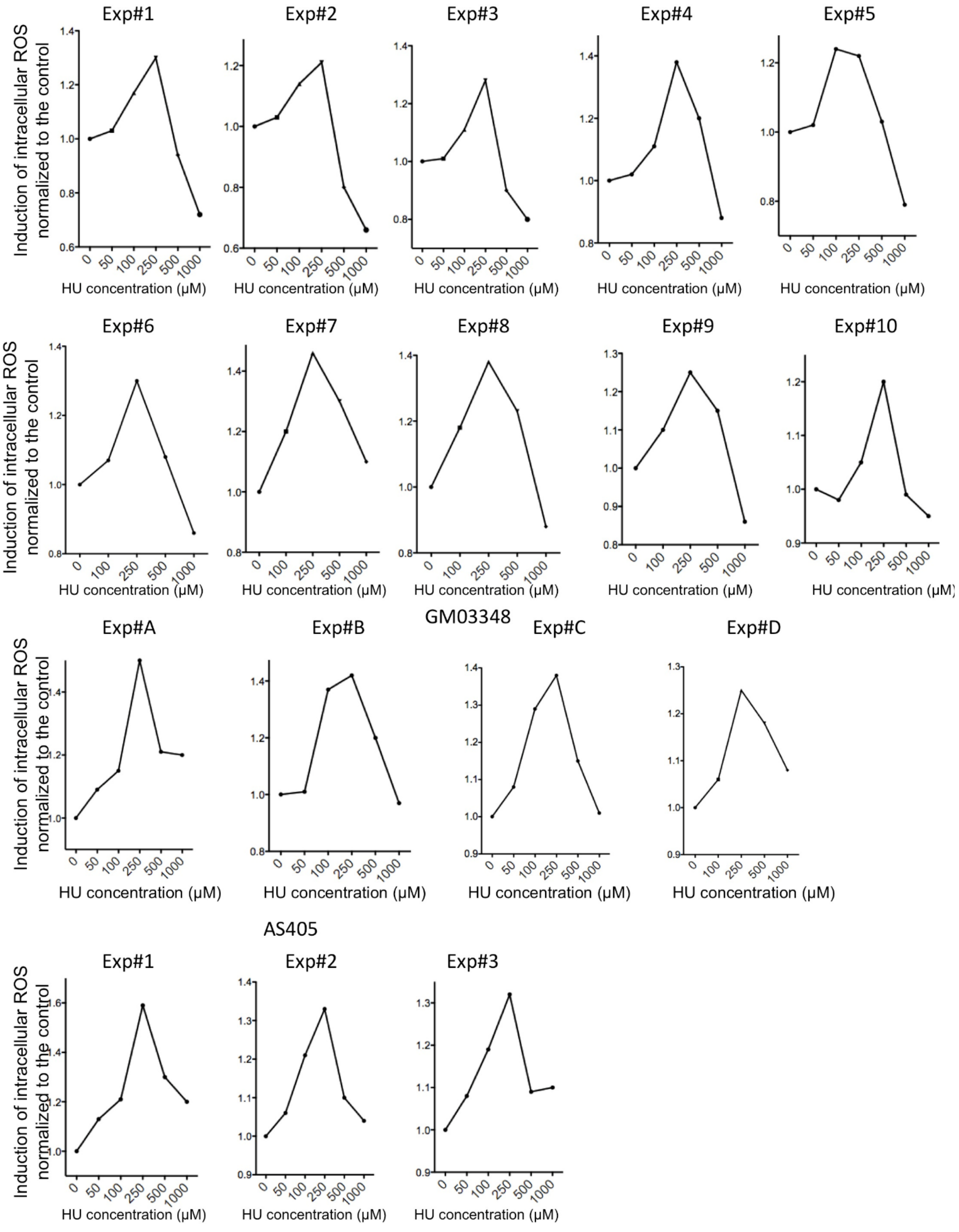

Supplementary data S2.

**Efficiency of silencing of DUOX1 and DUOX2 mRNA.**

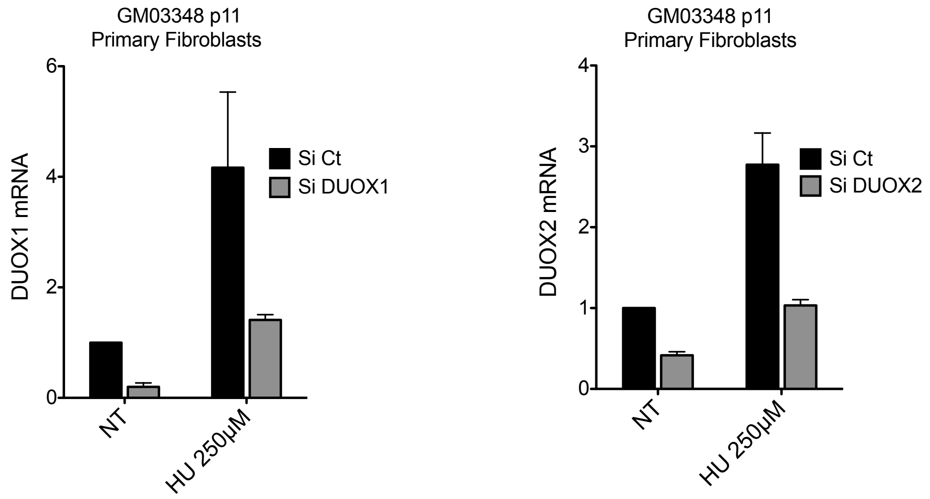


**Figure S2. Efficiency of silencing of DUOX1 and DUOX2 mRNA.**SiRNA against DUOX1 or DUOX2 or a control (scrambled siRNA) was transfected into primary fibroblasts, and HU was added or not. After 72 hrs, the cells were harvested, and DUOX1 and DUOX2 mRNA expression was analyzed by real-time qRT-PCR. The data represent the relative expression of DUOX mRNAs normalized to the control (scrambled siRNA) after the normalization of all samples to beta-actin mRNA expression.

**Supplementary data S3.**

#### Microarray analysis comparing primary human fibroblasts treated with HU anduntreated primary fibroblasts cells.

**Figure S3A.**

We performed a microarray analysis comparing primary human fibroblasts treated with HU250 µM for three days with non-treated cells. We used a cutoff of log2 (fold change - FC) > 0.5 and < −0.5, and an adjusted p-value of 0.05, 152 down- and 416 upregulated targets were found after HU treatment.

**Table S3A.1 : Microarray scores (see Excel table S3A.1)**
**Table S3A.2 :DAVID_Downregulated genes (see Excel table S3A.2)**
**Table S3A.3 : DAVID_Upregulated genes (see Excel table S3A.3)**

**Supplementary data S3B: Up-regulation of genes controlled by p65 (RelA) after HU.**

We searched for transcription factors that could regulate the expression of our microarray scores. The analysis revealed p65, a member of the NFκB signaling pathway, as a possible transcription factor activated after HU250 µM treatment

List of 69 NFκB65-controlled genes up-regulated after HU

NOG, SNORA12, IL4I1, GRIN3B, CPEB1, GDNF, GSTM5, IL11, LNX1, ARHGAP22, TRIM47, SLC24A3, SPINT2, IL1B, NOS3, NRG1, C17ORF96, KDELR3, H1F0, ICAM1, CAMK1G, EFNB1, NUDT14, RELB, TNFRSF14, SNORD96B, SNORD96A, HMGA2, PKIA, UCN2, TLCD1, PTGDS, KRT17, BTG2, CD82, CTSH, CCL2, PFKFB4, NFKBIE, NFKBIA, ATP6V1G2, NFIX, GPR68, SFN, DENND2D, FAM46C, LAPTM5, TEK, NUMB, HAAO, OBSL1, NEDD4L, NDRG2, DCLK1, C17ORF82, MAFF, IL6, NFE2, EEF1A2, PODXL, BIRC3, COG4, LAMA4, GMFG, SVIL, NOTCH4, FAM43B, MAP6, CEND1

**Supplementary data S4.**

**Impact of PARP inhibitors on RIR production.**

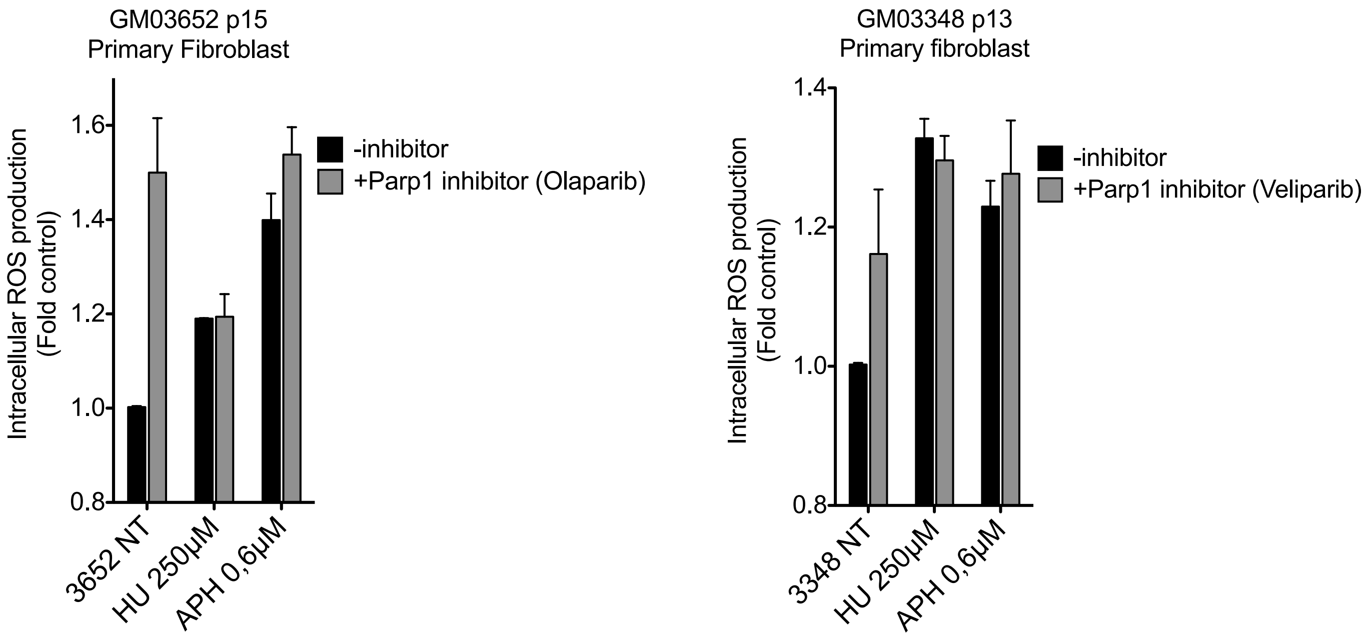


**Figure S4.Impact of PARP inhibitors on RIR production.**Effect of Olaparib and Veliparib (PARP1 inhibitor) on RIR production induced by hydroxyurea and aphidicolin intwo primary fibroblast strains.

**Supplementary data S5**

#### Whole genome sequencing

##### Mapping Statistics of whole genome sequencing

| Samples | # reads | Total mapped reads | % mapped read | Properly mapped reads | % Properly mapped reads | Duplication level |
|---|---|---|---|---|---|---|
| 3652_P16 Control | 112 935 167 | 112 624 848 | 99,73% | 107 887 102 | 95,53% | 20,16% |
| 3652_P16 HU 250 | 566 025 918 | 563 147 806 | 99,49% | 542 246 788 | 95,80% | 22,62% |
| 3652_P16 H2O2 75 | 405 439 770 | 404 476 760 | 99,76% | 393 296 262 | 97% | 22,83% |
| 3652_P16 HU250 + H2O2 75 | 214 247 944 | 213 563 714 | 99,68% | 206 964 192 | 96,60% | 22,34% |

**Supplementary data S6**

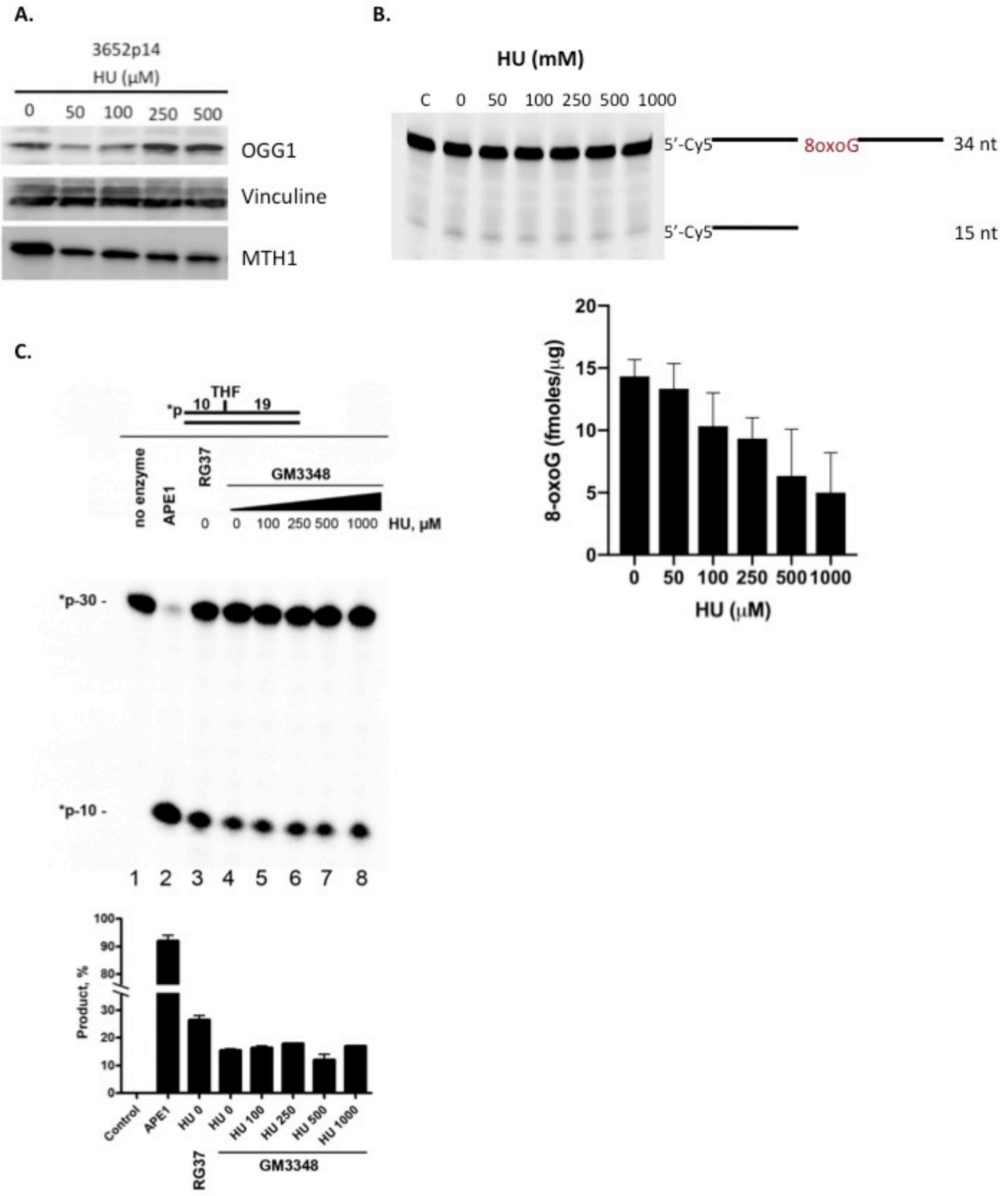


Figure S6. Impact of HU exposure on OGG1, MTH1 and APE1. A. Western blot of OGG1 and MTH1. B. **8-oxoG DNA glycosylase activity on whole cell extracts from cultures treated with HU.** 8-oxoG excision was determined on whole cell extracts by the cleavage assay on an oligonucleotide harbouring an 8-oxoG residue (upper panel). Values correspond to the mean three assays and their SEM(lower panel)**C. APE1 activity.**AP endonuclease activities in RG37 (SV40 trasnformed human fibroblast) and GM3348 (human primary fibroblasts) cell-free extracts after HU treatments. (Upper panel) PAGE analysis of AP endonuclease activities on double-stranded THF•C oligonucleotide substrate containing a single tetrahydrofuran (THF, a stable abasic site analogue). *p-10 band on the polyacrylamide gel corresponds to expected AP endonuclease activity. (Lower panel) Comparison of AP endonuclease activities in the experiments presented in upper panel . The error bars represent the standard deviation (n=3). For details, see the *Supplementary Materials and Methods.*

##### *Supplementary Materials and Methods S6*

**8-oxoG DNA glycosylase assay**. Cell pellets were sonicated in 20mM Tris-HCl, pH 7.5; 250mM NaCl; 1mM EDTA containing a cocktail of apoprotinin, antipain, and leupeptin (0,8$\mu$g/$\mu$l each). The homogenate was centrifuged at 20,000g for 30 min at 4°C and aliquots of the supernatant were stored at -80°C. Protein content was measured using a Bio-Rad assay kit (Bio-Rad Laboratories, Richmond, CA) with BSA as a standard. A 34-mer oligonucleotide containing an 8-oxoG at position 16 and labeled at the 5’ end with Cy5 was hybridized to its complementary oligonucleotide containing a cytosine opposite the lesion. In a standard reaction mixture protein extracts (20 or 10 $\mu$g in 10$\mu$l final volume) were added to a 20$\mu$l reaction mixture containing 150 fmoles of the 8-oxoG:C labeled duplex in 20mM Tris-HCl pH 7.1, 1mM EDTA, 200mM NaCl, 1mg/ml BSA and 5% glycerol. After 1h at 37°C, NaOH (0.1N final concentration) was added, and the mixture was further incubated for 15 min at 37°C and stopped by adding 4$\mu$l of formamide dye and heating for 5 min at 95°C. The products were resolved by denaturating7M urea -20% polyacrylamide gel electrophoresis. Gels were scanned and band intensities were quantified using a Typhoon PhosphoImager (GE Healthcare).

APE1 Activity assay.

***Preparation of cell extracts.*** Briefly, the cells pellet was washed in ice-cold PBS and incubated for 15 min at 4°C with shaking in 3 volumes of the lysis buffer containing 0.5 M KCl, 80 mM HEPES (pH 7.6), 0.1 mM EDTA, 2 mM DTT, 0.3% NP-40 and protease inhibitor cocktail (Complete EDTA-free, Roche). After centrifugation at 20000 rpm for 1 h at 4°C, the supernatants were collected and stored in 50% glycerol at -20°C for immediate use or at -80°C for longer storage. Protein concentration was determined by the Bradford assay.

***Oligodeoxyribonucleotide duplexes.*** Oligodeoxyribonucleotides were purchased from Eurogentec (Seraing, Belgium), including the following: d(TGACTGCATAXGCATGTAGACGATGTGCAT) 30mer, where X is either thymidine or tetrahydrofuranyl (THF), and complementary oligonucleotides. Oligonucleotide containing THF was 5'-end labeled by T4 polynucleotide kinase (New England Biolabs, OZYME France) in the presence of [$\gamma$-$^{32}$P]-ATP (3,000 Ci•mmol$^{-1}$) (PerkinElmer SAS, France) as recommended by the manufacturer. Complimentary oligonucleotides were annealed by heating at 65°C for 5 minutes and cooling slowly to room temperature for 1 h. The resulting duplexes are referred to as THF•C and T•A, containing either dC opposite the THF or dA opposite T, respectively.

***AP endonuclease assays.***The standard assay mixture for AP endonuclease activity (20 $\mu$l final volume) contained 0.2 pmol of the 5´-[$^{32}$P]-end labeled THF•C and 2 pmol of cold T•A oligonucleotide duplexes in 20 mMTris/HCl [pH 7.6], 50 mMKCl, 0.5 mM $MgCl_2$, 100 $\mu$g bovine serum albumin/ml, and either 12.5 $\mu$g/ml cell-free extract or 2.5 nM recombinant human APE1 enzyme (laboratory stock). Incubations were carried out at 37°C for 10 min. Desalted reaction products were heated at 65°C for 3 min and separated by electrophoresis in denaturing 20% (w/v) polyacrylamide gels (20:1, 7 M urea, 0.5xTBE). The gels were exposed to a Fuji FLA-3000 Phosphor Screen and analyzed using the ImageGauge V3.12 software.

**Supplementary data S7.**

**Table S7**: List of primersused for SYBR realtime RT-PCR.

| Gene Symbol | Forward primer 5' - 3' | Reverse primer 5' - 3' |
|---|---|---|
| SMC4 | GGCTGTATGGGCGAAAAAGAT | TTGTGGCTTGATCCAAGTTGT |
| LMB1 | GAAAAAGACAACTCTCGTCGCA | GTAAGCACTGATTTCCATGTCCA |
| MCM3 | TCAGAGAGATTACCTGGACTTCC | TCAGCCGGTATTGGTTGTCAC |
| HIST1H3F | TACTGTCGCCCTCCGTGAAA | CACCAGGTAAGCCTCGCAG |
| TOP2B | TTGGACAGCTTTTAACATCCAGT | GCACCATAACCATTACGACCAC |
| CCL2 | CAGCCAGATGCAATCAATGCC | TGGAATCCTGAACCCACTTCT |
| CXCL14 | CGCTACAGCGACGTGAAGAA | GTTCCAGGCGTTGTACCAC |
| CDKN1A (p21) | TGTCCGTCAGAACCCATGC | AAAGTCGAAGTTCCATCGCTC |
| IL4I1 | TGATGTCCGAGGATGGCTTCT | TGTACTGGAGTCTGTCGCTGA |
| CD82 | TGTCCTGCAAACCTCCTCCA | CCATGAGCATAGTGACTGCCC |

| | | |
|---|---|---|
| SOD2 | GCTCCGGTTTTGGGGTATCTG | GCGTTGATGTGAGGTTCCAG |
| IL6 | CCTGAACCTTCCAAAGATGGC | TTCACCAGGCAAGTCTCCTCA |
| GAPDH | ACAACTTTGGTATCGTGGAAGG | GCCATCACGCCACAGTTTC |
| ACTB | TGACCCAGATCATGTTTGAGA | TACGGCCAGAGGCGTACAGG |
